# CysQuant: simultaneous quantification of cysteine oxidation and protein abundance using data dependent or independent acquisition mass spectrometry

**DOI:** 10.1101/2023.07.22.550135

**Authors:** Jingjing Huang, An Staes, Francis Impens, Vadim Demichev, Frank Van Breusegem, Kris Gevaert, Patrick Willems

**Affiliations:** Department of Plant Biotechnology and Bioinformatics, Ghent University, Ghent, Belgium; Center for Plant Systems Biology, VIB, Ghent, Belgium; Department of Biomolecular Medicine, Ghent University, Ghent, Belgium; Center for Medical Biotechnology, VIB, Ghent, Belgium; VIB Proteomics Core, Ghent, Belgium; Department of Biochemistry, Charité - Universitätsmedizin Berlin, Berlin, Germany

## Abstract

Protein cysteinyl thiols are susceptible to reduction-oxidation reactions that can influence protein function. Accurate quantification of cysteine oxidation is therefore crucial for decoding protein redox regulation. Here, we present CysQuant, a novel approach for simultaneous quantification of cysteine oxidation degrees and protein abundancies. CysQuant involves light/heavy iodoacetamide isotopologues for differential labeling of reduced and reversibly oxidized cysteines analyzed by data-dependent acquisition (DDA) or data-independent acquisition mass spectrometry (DIA-MS). Using plexDIA with *in silico* predicted spectral libraries, we quantified an average of 18% cysteine oxidation in *Arabidopsis thaliana* by DIA-MS, including a subset of highly oxidized cysteines forming disulfide bridges in AlphaFold2 predicted structures. Applying CysQuant to Arabidopsis seedlings exposed to excessive light, we successfully quantified the well-established increased reduction of Calvin-Benson cycle enzymes and discovered yet uncharacterized redox-sensitive disulfides in chloroplastic enzymes. Overall, CysQuant is a highly versatile tool for assessing the cysteine modification status that can be widely applied across various mass spectrometry platforms and organisms.

## Introduction

Protein post-translational modifications (PTMs) diversify the complexity and functionality of the proteome. Cysteine (Cys) is amongst the most reactive amino acids, whose nucleophilic thiol side-chain (–RSH) is susceptible to oxidative modifications. Cys thiols react with reactive oxygen species such as hydrogen peroxide (H_2_O_2_) to form sulfenic acid (–RSOH), an unstable intermediate that can react with other reactive sulfur species. These reactions include formation of disulfides (–RSSR–) with protein thiols, *S*-glutathione adducts (–RSSG) with glutathione, or persulfides (–RSSH) with hydrogen sulfide (*1*). Cys oxidation can be enzymatically catalyzed, such as by protein disulfide isomerases that introduce disulfide bridges during protein folding (*2*), or indirectly by thiol peroxidases under specific conditions through disulfide exchange (*3–5*). Such cysteine oxidations can be reverted (reduced) by a group of redox enzymes including thioredoxins and glutaredoxins, collectively referred to as redoxins. These redoxins convey reducing power from redox metabolites such as glutathione, NAPDH and ascorbate (*6, 7*) and can protect protein thiols from overoxidation to sulfinic and sulfonic acids (–RSO_2_H and –RSO_3_H, respectively), which may lead to protein inactivation.

Diverse mass spectrometry (MS) based proteomics methods have been developed to profile protein Cys oxidation (*1*). Such methods can be directed towards specific oxidative Cys modifications via chemoselective probes (*8–10*) or trapping substrates of redoxins (or other redox-active enzymes) via inter-protein disulfides (*11, 12*). Additionally, more commonly used workflows make use of reducing agents, typically dithiothreitol (DTT) or tris(2-carboxyethyl) phosphine (TCEP), to indiscriminately reduce oxidized Cys and afterwards record these reversible oxidative modifications indirectly. Such workflows typically consist of four main steps: (*i*) blocking native, reduced thiols, (*ii*) reduction of reversible Cys oxidative modifications by DTT/TCEP (or other reducing agents), (*iii*) labelling of newly reduced thiols, and (*iv*) quantitative MS analysis. Moreover, by using chemically distinguishable mass tags that label Cys thiols prior or after reduction (step *iii*), an absolute ratio or degree (%) of protein Cys oxidation can be calculated. While reporting relative levels of Cys oxidative modifications useful, it is of higher value to calculate degrees (or occupancy, stoichiometry) of Cys oxidation (*13, 14*). The OxiCAT method first introduced the concept of Cys occupancy analyses using isotope coded affinity tag reagents (*15*), while later, other isotopologous and isobaric Cys reagents have been adopted to quantify the degree of Cys oxidation (*16–18*). However, these methods remain scarce and would benefit from a more widely adoptable workflow.

In the past years, advanced MS instruments and acquisition methods have allowed to obtain an increasingly deeper proteome coverage, data completeness, throughput and sensitivity (*19–21*). Here, data-independent acquisition (DIA) in particular has been a major facilitator. Indeed, in DIA mode, fragmentation of multiple peptide precursor ions is performed in predetermined, consecutive *m/z* windows over time, thereby providing a more reproducible and completer identification than classical data-dependent acquisition (DDA) workflow (*22*). DIA-MS has also been used in thiol-based redox proteomic workflows for quantifying the intensity of different Cys alkylation reagents (*23*) and for reactivity profiling and ligand screening (*24*).

Given that DIA-MS analysis allows for multiplexed non-isobaric quantification (*21*), we developed a novel labeled Cys redox proteomic method using light and heavy iodoacetamide isotopologues, named ‘CysQuant’. CysQuant allows simultaneously quantifies the degree of Cys oxidation and protein abundance using both DIA-MS and traditional DDA-MS. As a proof-of-concept, we applied CysQuant to study the response of proteome Cys thiols to excess light in *Arabidopsis thaliana* seedlings. This efficiently quantified the reduction of characterized regulatory disulfides in the Calvin-Benson cycle and revealed novel redox-sensitive disulfides in plastidial enzymes. With deep learning predictors trained on iodoacetamide-modified peptides, the CysQuant workflow is fully compatible with *in silico* predicted spectral libraries in a DIA-only analysis with multiplexed quantification using the plexDIA module of DIA-NN (*21, 25*). Altogether, CysQuant is a highly reliable and powerful tool for Cys oxidation quantification in redox biology, compatible with a wide range of mass spectrometry platforms.

## Results

### CysQuant workflow: simultaneous quantification of the Cys oxidation degree and protein abundance

CysQuant uses an isotopic labelling strategy that ultimately leads to simultaneous quantification of the proteome and profiling of Cys oxidation degrees (Fig. 1). In the CysQuant workflow, plant samples were grinded to fine powder in liquid nitrogen and immediately transferred into a trichloroacetic acid (TCA) buffer to render reduced thiols protonated and non-reactive. Following protein precipitation, reduced Cys thiols were labelled with light iodoacetamide (IAM^0^, C_2_H_4_INO, L) in a strong denaturing buffer. Further protein processing steps proceeded using an adapted version of the suspension trapping (S-Trap) protocol, a fast and efficient sample preparation workflow (*26*) that can be performed under strongly denaturing conditions. In brief, proteins were captured on the S-Trap column, on which oxidized Cys thiols were reduced with TCEP and labelled with heavy IAM (IAM^4^, ^13^C_2_D_2_H_2_INO, H). After trypsin digestion, differentially labelled Cys peptides and unmodified peptides lacking Cys were analyzed by liquid chromatography coupled to tandem mass spectrometry (LC-MS/MS) in both data-dependent acquisition (DDA) as well as data-independent acquisition (DIA) modes. Intensities of IAM^0^- and IAM^4^-modified cysteine-containing peptides (Δ 4 Da) were quantified using computational algorithms typically used for other non-isobaric labelling reagents SILAC, mTRAQ, though now with light/heavy (L/H) ratios reflecting the degree of Cys oxidation. Label-free quantification of unlabeled, non-cysteine peptides enables to simultaneously compare protein abundance across samples. For DDA analysis, IAM^0^ and IAM^4^ were specified as fixed Cys labels in MaxQuant (*27–29*). In DIA analysis, we used the plexDIA module of DIA-NN to quantify L/H Cys peptide ratios (*21, 25*). Aside from MS2 fragment ion quantification, accurate MS1 precursor ion-based quantification is obtained with plexDIA (*21*). Importantly, this represents a DIA-only workflow, independent from DDA analysis, using *in silico* spectral libraries generated by DIA-NN that can be readily used for labelled Cys quantification as the deep learning predictor is trained on IAM-modified Cys peptides.

**Fig. 1.**
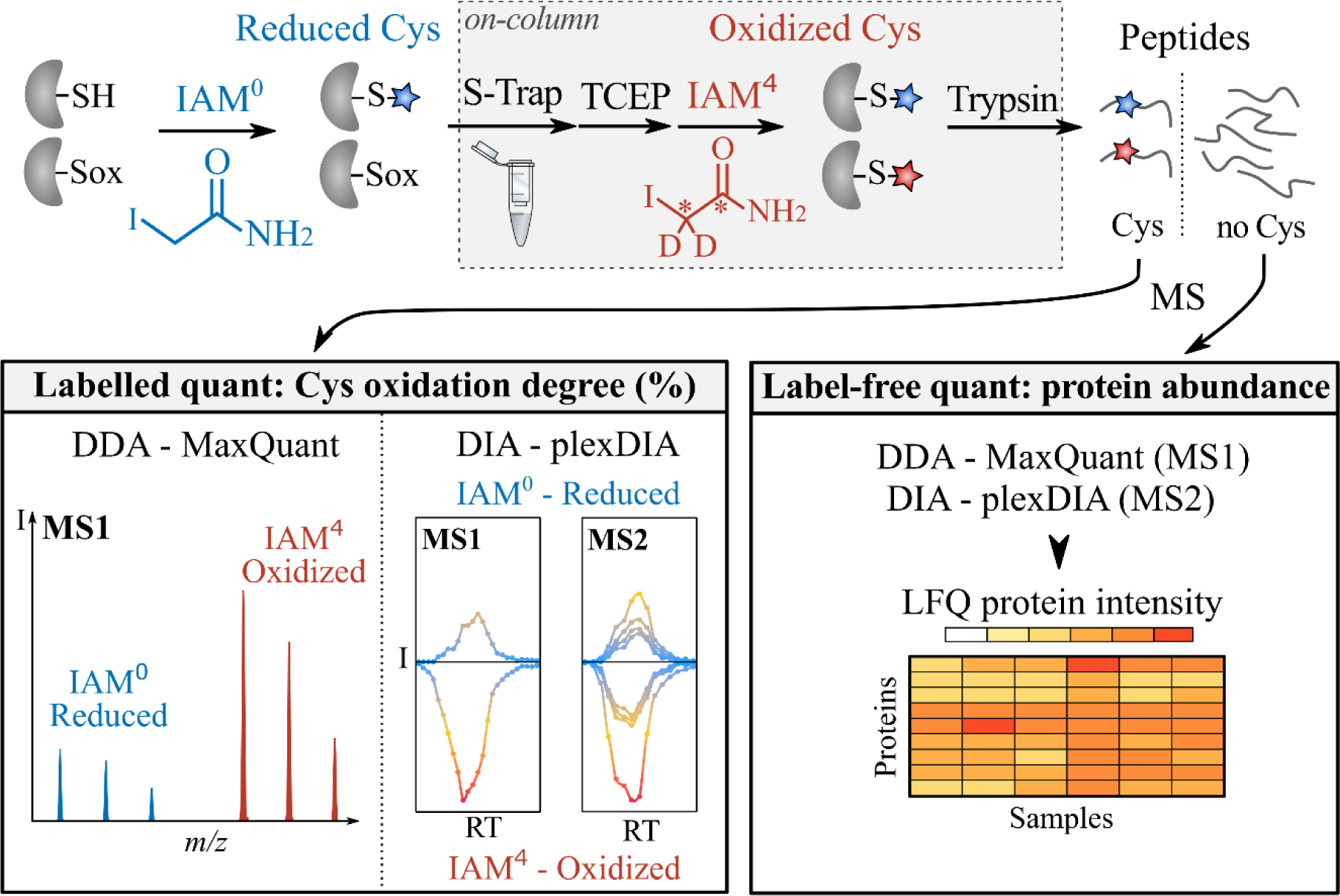
CysQuant workflow for cysteine oxidation degree and protein abundance quantification. Following protein extraction in an acidic buffer to protonate and inactive reduced Cys thiols, these reduced thiols are labelled with (light) IAM^0^ (^12^C2H4INO, blue). Then, proteins are processed using an adapted version of the suspension trapping (S-Trap) protocol (*26*). After protein loading on the S-Trap column, oxidized Cys thiols are reduced with tris(2-carboxyethyl)phosphine (TCEP) and labelled with heavy IAM^4^ (^13^C2D2H2INO, red). After trypsin digestion, Cys and non-Cys peptides enable labelled and label-free quantification to determine the Cys oxidation degree and protein abundance within and across samples, respectively. Mass spectrometry (MS) analysis was performed for each sample in parallel by data-dependent acquisition (DDA) analyzed by the MaxQuant software suite, and by data-independent acquisition, analyzed by the plexDIA module of DIA-NN for labelled and label-free analysis respectively.

### Profiling the redox proteome under excess light in *Arabidopsis thaliana*

We used the CysQuant workflow to study the excess light (EL) response in *Arabidopsis thaliana* (Arabidopsis) seedlings. Light is intimately linked with redox signaling in plants as photosynthetic electron transport in the chloroplast converts light to reducing power that can be transduced to Cys thiols of target proteins via redoxin enzymes (*30, 31*). EL causes photoinhibition, impeding PSII photochemistry that can be measured by altered photosynthetic parameters such as a decrease in maximum photosystem II quantum efficiency (F_v_’/F_m_’) (*32*). Exposing 23-day-old seedlings from low light growth conditions (50 µE m^−2^ s^−1^) to twenty-fold more intense light exposure (1,000 µE m^−2^ s^−1^) resulted in a reversible photoinhibition evidenced by a transient F_v_’/F_m_’ decrease (Fig. 2A, Fig. S1). Here, we applied the CysQuant workflow to quantify cysteine oxidation levels and protein abundance of above-ground seedling tissues prior to the EL treatment and 10, 60 or 180 min after EL exposure in four biological replicates.

**Fig. 2.**
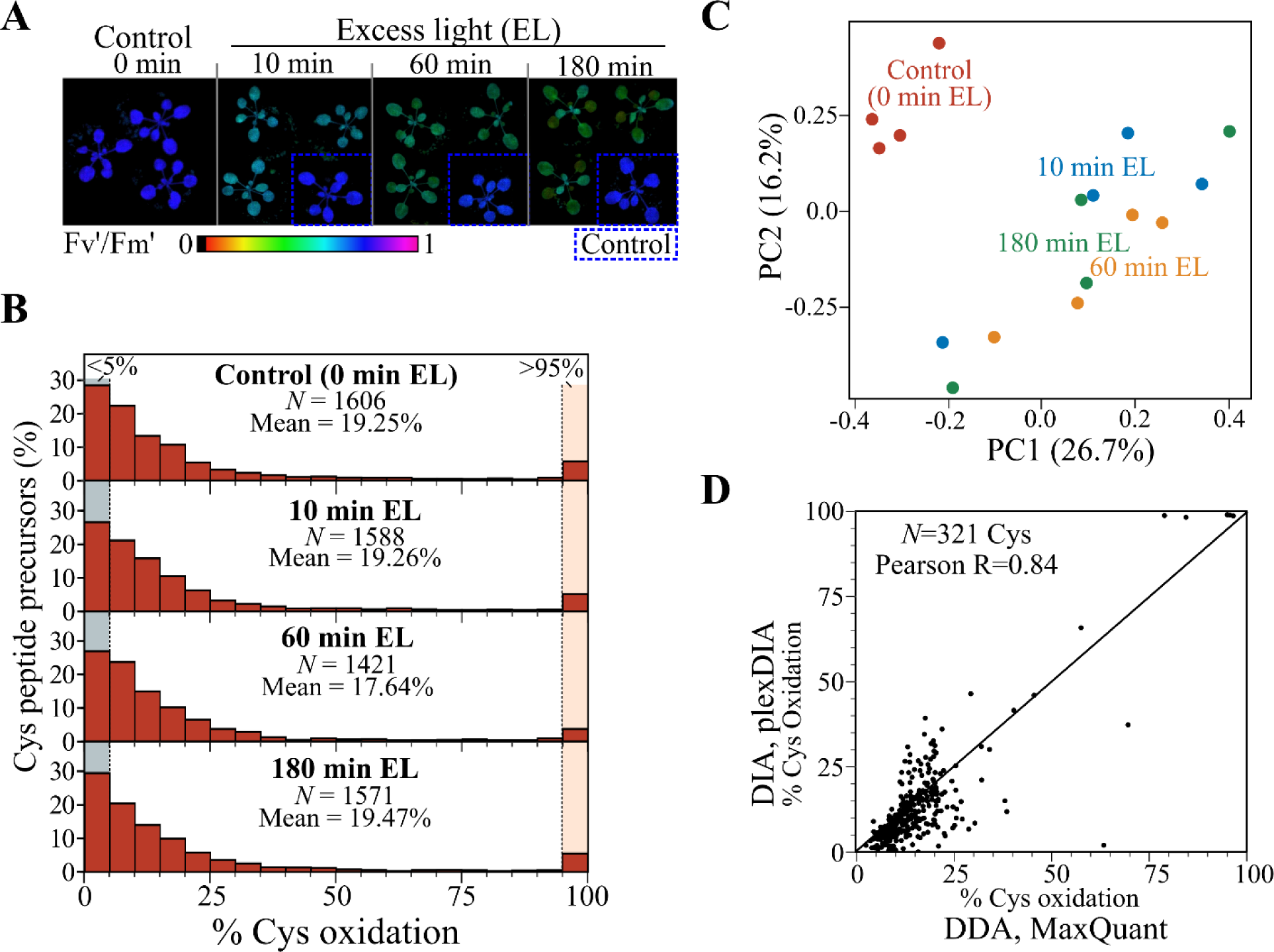
CysQuant profiling cysteine oxidation changes in Arabidopsis in response to excess light. (**A**) Representative pictures of 23-day-old *Arabidopsis thaliana* seedlings transferred from low light growth conditions to excess light (EL, 50 to 1000 µE m^−2^ s^−1^), displaying a decrease in maximum photosystem II quantum efficiency (Fv’/Fm’) (*N*=3). Control seedlings not exposed to EL are indicated in blue dotted boxes. For CysQuant analysis, seedlings were collected at 0, 10, 60 or 180 minutes of EL stress. (**B**) Distribution of Cys oxidation degree (%) quantified by plexDIA in at least three out of four replicates per condition. Histogram boxes of lowly (< 5%) and highly oxidized Cys (> 95%) were displayed in a grey and orange background, respectively. For a detailed overview see Data S1A. (**C**) Principal component analysis (PCA) of samples using plexDIA quantified oxidation degree of 516 Cys quantified in all samples. PCA was performed using the *prcomp* function in R. (**D**) Correlation of the oxidation degree (%) of 321 Cys quantified by plexDIA and MaxQuant in all four replicates of the control condition (0 minutes of EL stress). For other conditions see Fig. S3.

A MaxQuant label-free search, with IAM^0^ and IAM^4^ specified as variable modifications, identified ∼96-98% of spectra matching Cys peptides labelled with IAM^0^ (reduced Cys) and ∼2 to 4% with IAM^4^ (oxidized Cys, Fig. S2), thus verifying efficient IAM labelling in these samples and suggesting an overall reduced Cys state. This overall low degree of Cys oxidation was also evident upon quantification by plexDIA, averaging ∼18 to 19% of Cys oxidation across all conditions (Fig. 2B, Data S1A). In each condition, there is a clear prevalence of strongly reduced Cys (less than 5% oxidized), for instance accounting for 457/1606 Cys peptide precursors (28.5%) in the control condition (Fig. 2B) (Data S1A). Noteworthy, this includes 122 Cys peptide precursors only present in the IAM^0^ (reduced) channel, interpreted here as a fully reduced cysteines. On the other hand, strongly oxidized Cys (over 95%) were also found in each condition, accounting for 91 Cys in the control condition, of which 24 were exclusively present in the IAM^4^ (oxidized) channel and thus fully oxidized. While EL did not cause a major effect on the overall Cys oxidation degree distribution, principal component analysis (PCA) clearly shows separation of the control and EL treated samples (Fig. 2C).

MaxQuant analysis verified the overall reduced state of the analyzed proteomes with average Cys oxidation degrees of ∼18 to 19%, though outputting lower numbers of strongly reduced (< 5%) or oxidized Cys (> 95%) (Fig. S3, Data S1B). Notably, Cys oxidation degrees quantified by both MaxQuant and plexDIA in all replicates show a high correlation per condition (average r = 0.815) (Fig. 2D), with a median coefficient of variation (CV%) averaging 10.1% and 13.2% in plexDIA and MaxQuant analysis, respectively (Fig. S4). This confirms CysQuant to be compatible with both DDA and DIA platforms. Lastly, we applied CysQuant to non-EL exposed seedlings using a (non-acidic) methanol-chloroform protein extraction (Data S2). This resulted in a nearly 20% increase in Cys oxidation degree (average 39.5%, Fig. S5), emphasizing the importance of acidic protein extraction to render free thiols protonated during initial sample processing.

### Cysteines part of disulfides or secreted proteins are strongly oxidized

Protein disulfides are inaccessible to the initial IAM^0^ proteome labeling, but they can be reduced and labeled with IAM^4^ in subsequent step. We anticipated structural disulfides to corresponds to the observed subset of strongly oxidized Cys (Fig. 2B). To obtain a proteome-wide proxy of such structural disulfides, we parsed Cys sulfur atom pairs within 2.5 Å distance in AlphaFold2 predicted plant protein structures (*33*). Cysteines that are part of disulfides in predicted protein structures presented an average oxidation degree of 82.5% across all conditions, in contrast to other Cys that were on average 13.6% oxidized (Fig. 3A). Moreover, several peptides carrying such disulfide Cys were only identified in the IAM^4^ channel and were thus interpreted as being fully oxidized. Taking a closer look to 76 Cys-containing peptides that were only present in the IAM^4^ channel in at least four out of sixteen samples, reveals these peptides to be often quantified to be over 90% oxidized in other samples (Fig. 3B), further supporting their strongly oxidized state. Moreover, 68 of these 76 Cys (89.5%) were part of AlphaFold2 predicted disulfides.

**Fig. 3.**
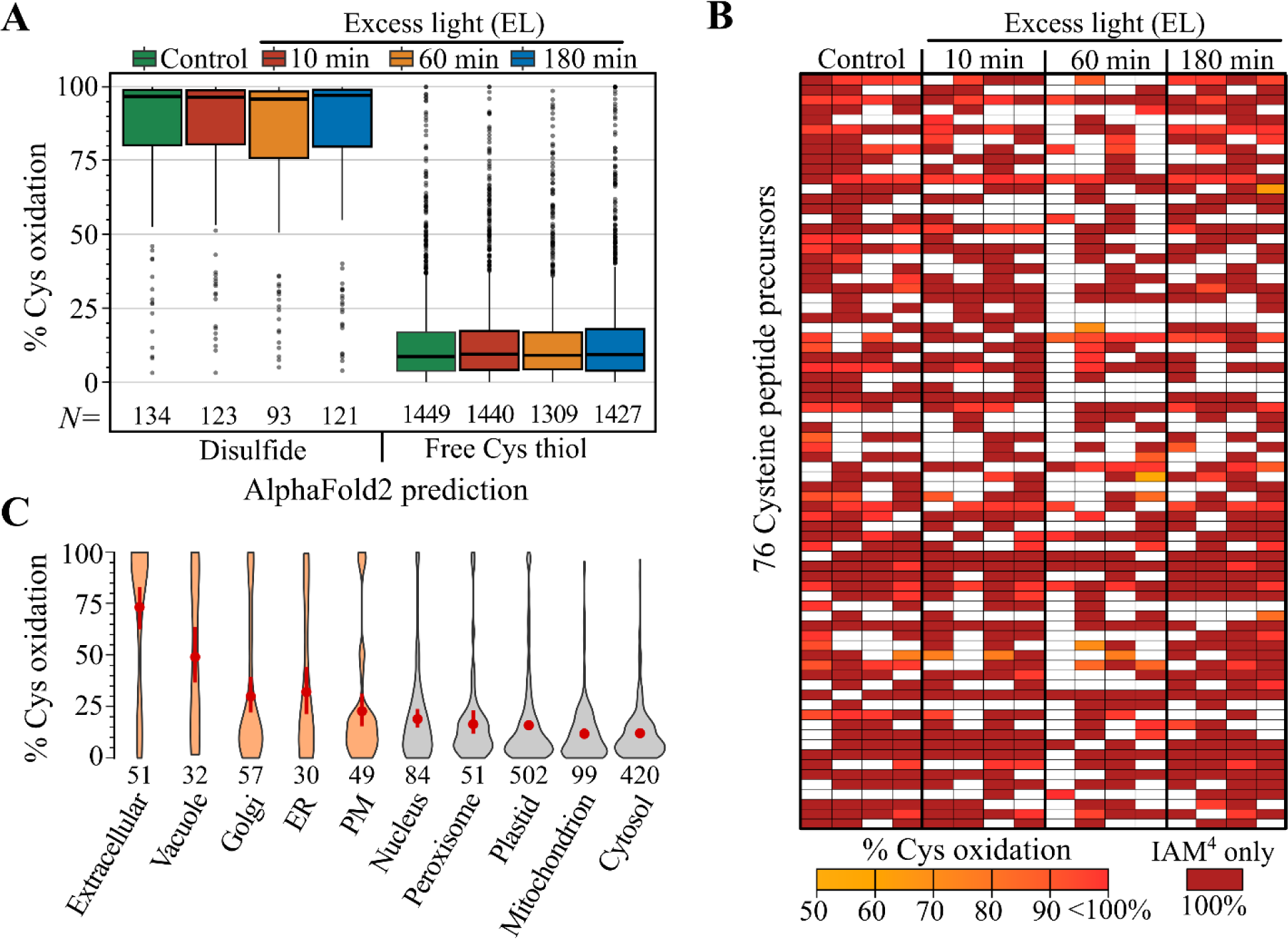
Cysteines that are part of disulfides or found in extracellular proteins are highly oxidized. (**A**) Oxidation degree for Cys quantified in three out of four replicates categorized according the AlphaFold2-based prediction as being part of a disulfide or rather a free thiol (*33*). For metal ion predictions see Fig. S6A. (**B**) Heatmap displaying the oxidation degree for 76 Cys with at least four IAM^4^-only channel quantifications. Missing values are colored white. (**C**) Oxidation degree for Cys quantified in three out of four replicates in the control condition according to the subcellular locations predicted by the SUBAcon algorithm (*75, 76*). Compartments associated with the secretory pathway are colored in orange, and the mean oxidation degree ± sd is indicated in red with the number of Cys peptide precursors displayed under the violin plot. For plots of excess light conditions see Fig. S6B.

Besides disulfides, metal ion coordination by Cys has been predicted for the entire *Arabidopsis* proteome by making use of AlphaFold2 predicted protein structures (*33, 34*). However, the performed low pH protein extraction would destabilize metal ion-binding pockets, causing Cys involved in metal ion binding to not exhibit higher oxidation degrees in our analysis (Fig. S6A). In contrast, the non-acidic methanol-chloroform extraction method maintained metal ion coordination, as relatively higher oxidation degrees for metal-binding Cys were identified (Fig. S6A). Finally, the majority of Cys thiols not participating in disulfides or metal ion coordination in AlphaFold2 predicted structures were on average 13% (Data S1A). However, a subset of Cys displayed higher oxidation degrees that might reflect a structural property or a functional activity. For instance, the active sites Cys111 and Cys41 of GLUTATHIONE PEROXIDASE 1 and 2 displayed 58.2% and 54.1% oxidation degrees in control conditions, respectively, in line with their H_2_O_2_ detoxification catalysis.

Lastly, we evaluated Cys oxidation degrees in terms of the subcellular location of their respective proteins. Extracellular proteins and proteins predicted to reside in other compartments associated with the secretory pathway such as the vacuole, Golgi apparatus, ER and plasma membrane (PM) presented the highest average oxidation degree, while more reduced Cys states were observed for chloroplast, mitochondrial and cytosolic proteins (Fig. 3C, Fig. S6B). This observation is consistent with the higher occurrence of AlphaFold2 predicted disulfides in the former compartments (*33*). In conclusion, quantifying steady state Cys oxidation levels can contribute to functional annotation of Cys.

### Reduction of disulfides in chloroplast carbon assimilation pathways under excess light

To identify EL-induced changes in Cys oxidation degrees, we performed pairwise differential statistics for each EL timepoint against the control condition (0 min EL) for Cys quantified in at least three out of four replicates. We detected for 23, 35 and 26 Cys after 10, 60 and 180 min of EL treatment, respectively, significant changes (Welch’s t-test *P* < 0.05) of an at least 5% absolute change in oxidation (Fig. 4A, Data S1). After 10 min of EL, more than half of these Cys (12/23) were found in chloroplastic proteins – of which eight Cys were significantly reduced and predicted as disulfides (Fig. 4A – triangles). Some of these corresponded to regulatory disulfides of Calvin-Benson enzymes and accessory proteins, which are known to undergo light-dependent reduction by the ferredoxin/thioredoxin (Fd-Trx) redox relay (*30, 35*) (Fig. 4b). For instance, Cys constituting the regulatory disulfide Cys451-Cys471 in the longer α-isoform of RUBISCO ACTIVASE (RCA-α) were found to be approximately 38 to 45% more reduced under EL (Fig. 4C-D). Likewise, rapid and consistent reduction of disulfide Cys was observed for the Calvin-Benson cycle proteins PHOSPHORIBULOKINASE (PRK), CALVIN CYCLE PROTEIN CP12-2 and GLYCERALDEHYDE-3-PHOSPHATE DEHYDROGENASE GAPB (Fig. 4C-D). The MaxQuant DDA analysis verified the reduction of PRK Cys295 and GAPB Cys434 by EL in three out of four replicates (Fig. S7), while Cys of CP12-2 and RCA-α lacked comprehensive quantification across replicates (Data S1B).

**Fig. 4.**
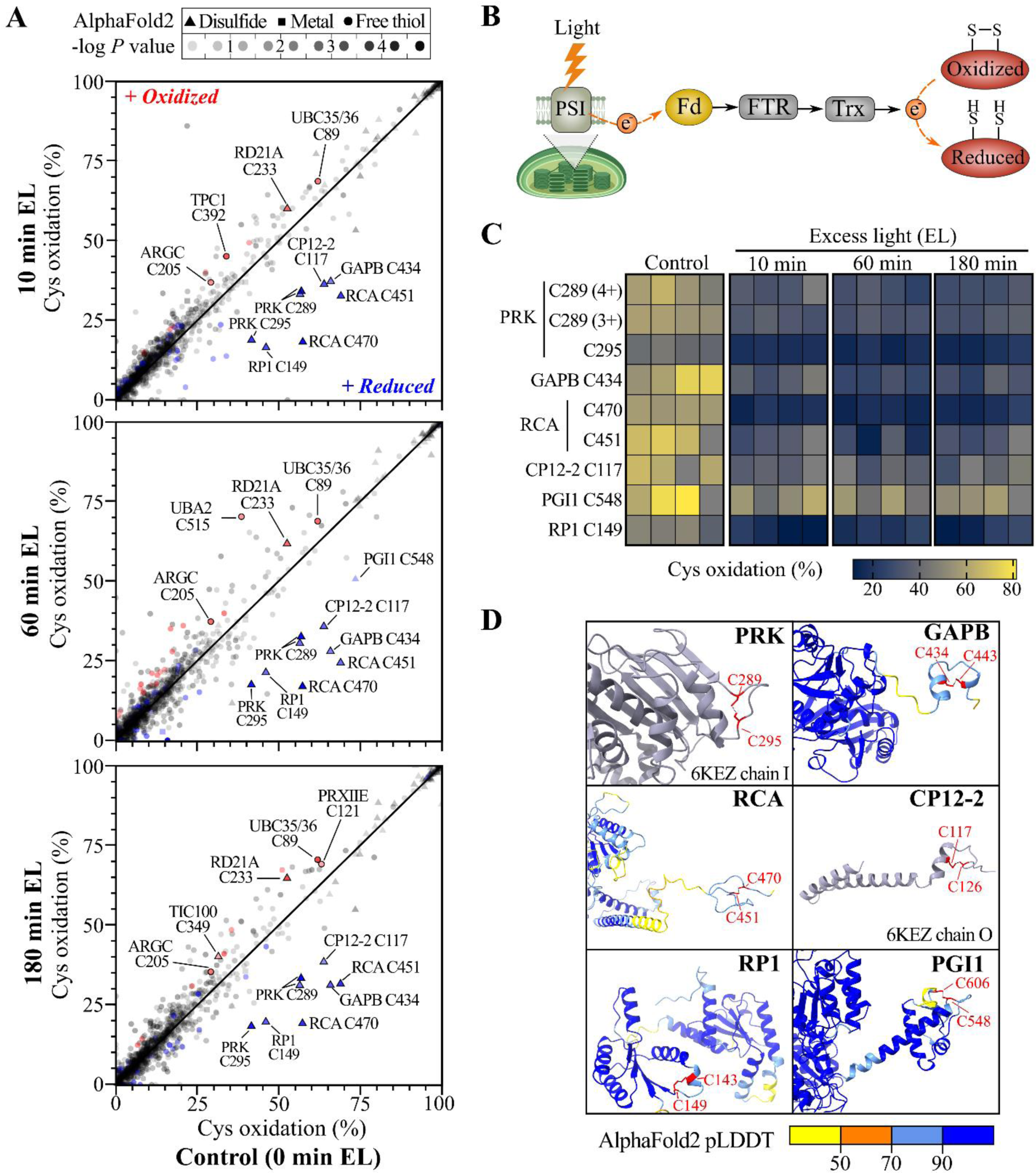
Reduction of disulfides in carbon metabolic enzymes in the chloroplast by excess light. (**A**) Scatterplot of the degree of Cys oxidation in unstressed plant leaves (control, x-axis) and leaves exposed to 10, 60 or 180 minutes of excess light (EL, y-axis, *top to bottom*). Dot transparency is according the *P* value of the Welch’s t-test (EL/control), with significantly altered oxidation (*P* < 0.05, ≥ 5% change in the degree of oxidation) indicated in red (more oxidized) or blue (more reduced). Shapes are according AlphaFold2 predicted Cys annotation (*33*). (**B**) Simplified scheme of chloroplast light redox regulation. Ferredoxin (Fd) is reduced by the photosynthetic electron transfer chain at photosystem I (PSI). Fd-Trx reductase (FTR) transfers electrons (e^−^) via disulfide relays from Fd to Trxs that can reduce target proteins in the chloroplast (*30*). Figure made using BioRender.com. (**C**) Heatmap of the degree of Cys oxidation (%) for significantly altered Cys disulfides in chloroplast proteins. (**D**) Chloroplast protein structures that had (predicted) disulfides reduced under EL (red). Structures were derived from crystal structures (*61*) or AlphaFold2 predictions (*51*) and visualized using ChimeraX (v1.6.1) (*77*). AlphaFold2 residues are colored by per-residue confidence score (pLDDT) with > 90 being highly confident and < 50 of very low confidence.

Aside from these well-characterized regulatory disulfides (*36*), we identified additional redox-sensitive chloroplast proteins related to carbon assimilation pathways. For example, the Cys548 of PHOSPHOGLUCOSE ISOMERASE 1 (PGI1) showed a significant 22.9% reduction (Welch’s t-test *P 0.036*) in Cys oxidation degree after 60 minutes EL, and a 20.3% reduction after 10 minutes EL that just above the significance threshold (Welch’s t-test *P 0.052*). AlphaFold2 prediction revealed PGI1 Cys548 to be in close proximity to Cys606 in an extended C-terminal domain (Fig. 4C-D), which resemble the redox-regulated C-terminal domains of RCA-α and GAPB. Cys149 of the chloroplastic PPDK REGULATORY PROTEIN 1 (RP1) was also significantly (∼25%) reduced in all EL timepoints (Fig. 4C-D). Lastly, MaxQuant consistently identified ∼15% reduction in Cys oxidation degrees of POST-ILLUMINATION CHLOROPHYLL FLUORESCENCE INCREASE (PIFI) Cys124 in two out of four replicates. Notably, Cys 124 forms a disulfide with Cys203 in the AlphaFold2 predicted PIFI protein structure (Fig. S7D). PIFI is involved in nonphotochemical reduction of the plastoquinone pool, and an electron carrier activity was attributed to two conserved Cys in its C-terminal domain (*37*).

Besides EL-induced reduction of Cys, we also observed increased oxidation. For instance, the Cys233 of papain-like Cys proteinase RD21A showed 7.5%, 9.2%, and 12.2% increased oxidation after 10, 60 and 180 minutes of EL, respectively, and forms a disulfide bond with Cys192 in the AlphaFold2 predicted structure. Induced oxidation in all EL timepoints (+6 to 8%) was also observed for the active site Cys205 of a putative N-acetyl-gamma-glutamyl-phosphate reductase in the chloroplast (ARGC) and the catalytic Cys89 residues of the E2 ubiquitin-conjugating enzymes UBC35/36 (Data S1A). The increased Cys oxidation for both enzymes could potentially relate to differential enzymatic activity, such as increased thioester formation for UBC35/36. Cys of other ubiquitin-system related enzymes such as Cys515 of ubiquitin-activating enzyme UBA2 and Cys2843 of E3 ubiquitin ligase UPL2 exhibited increased oxidation levels after 60 and 180 minutes of EL, respectively. Additionally, we noticed a rapid increase in oxidation (+ 11.2%) of Cys392 in the second Ca^2+^ binding EF-hand domain of the Ca^2+^-channel TWO-PORE CHANNEL 1 (TPC1) after 10 minutes EL for Cys392. Conversely, Cys oxidation was increased for certain proteins at later time point (180 minutes) of EL treatment. For example, the active site Cys121 of choloroplastic PEROXIREDOXIN IIE (PRXIIE) exhibited 6.2% increased oxidation, suggesting an increased H_2_O_2_ detoxification activity during prolonged EL exposure. Furthermore, we observed an increased oxidation after 180 min of EL for potential disulfide-containing Cys349 in TRANSLOCONS AT THE INNER ENVELOPE MEMBRANE OF CHLOROPLASTS 100 (TIC100), which facilitates the import of nuclear-encoded chloroplast proteins from the cytosol.

### Differential protein expression under excess light points to photoprotective mechanisms

As no enrichment step for Cys-containing peptides is incorporated in the CysQuant workflow (Fig. 1), peptides without Cys are amply recorded and can be used to evaluate protein abundance. In the MaxQuant label-free analysis, with IAM^0^ and IAM^4^ as variable modifications, 36,313 non-Cys peptides were identified, while plexDIA identified 51,540 non-Cys peptides. For differential protein abundance analysis, MS1 level (MaxQuant) and MS2 level (plexDIA) quantification features of non-Cys peptides were used by MSstats (*38*) (Data S3A and S3B, respectively). The plexDIA analysis revealed an increasing number of altered protein levels over time, with 33, 87 and 93 differentially abundant proteins (*P* ≤ 0.01 and absolute fold change > 1.5) after 10, 60 and 180 minutes of EL treatment, respectively (Fig. 5A). More specifically, after 60 minutes of EL, there was a strong decrease in the abundance of 72 proteins, while only 15 proteins were more abundant. In contrast, following 180 minutes of EL treatment, 57 proteins showed increased abundance while the abundance of 36 proteins decreased. Among the proteins that showed the most significant changes in abundance was GRANULE-BOUND STARCH SYNTHASE 1 (GBSS1), the main enzyme synthesizing amylose in starch granules (*39*), here found to be ∼3.65-fold downregulated after 60 and 180 minutes of EL treatment (Fig. 5B). In addition, the abundance of uncharacterized protein AT5G35970.1 and DORMANCY-ASSOCIATED GENE 2 (DRM2) were significantly repressed after prolonged EL exposure. Another enzyme that showed significant changes in abundance was CHLOROPHYLL A OXYGENASE (CAO), which oxygenates chlorophyll (Chl) *a* to Chl *b* in the chloroplast. We observed a 1.58-fold decrease in protein levels after 180 minutes of EL, which is consistent with previous reports of lower CAO protein levels and altered Chl levels upon EL treatment (*40*). We also considered proteins that were only detected by LC-MS/MS either in three replicates of the control condition or the EL condition (Data S3). This revealed the photoprotective thylakoid protein EARLY LIGHT-INDUCED PROTEIN 1 (ELIP1) that influences Chl levels (*41–43*), was identified only after 60 and 180 minutes of EL stress (Fig. 5B). Notably, ELIP1 is a classical marker of EL stress in plants that is undetectable under low light conditions (*42*), thus matching with the anticipated pattern in our experimental set-up.

**Fig. 5.**
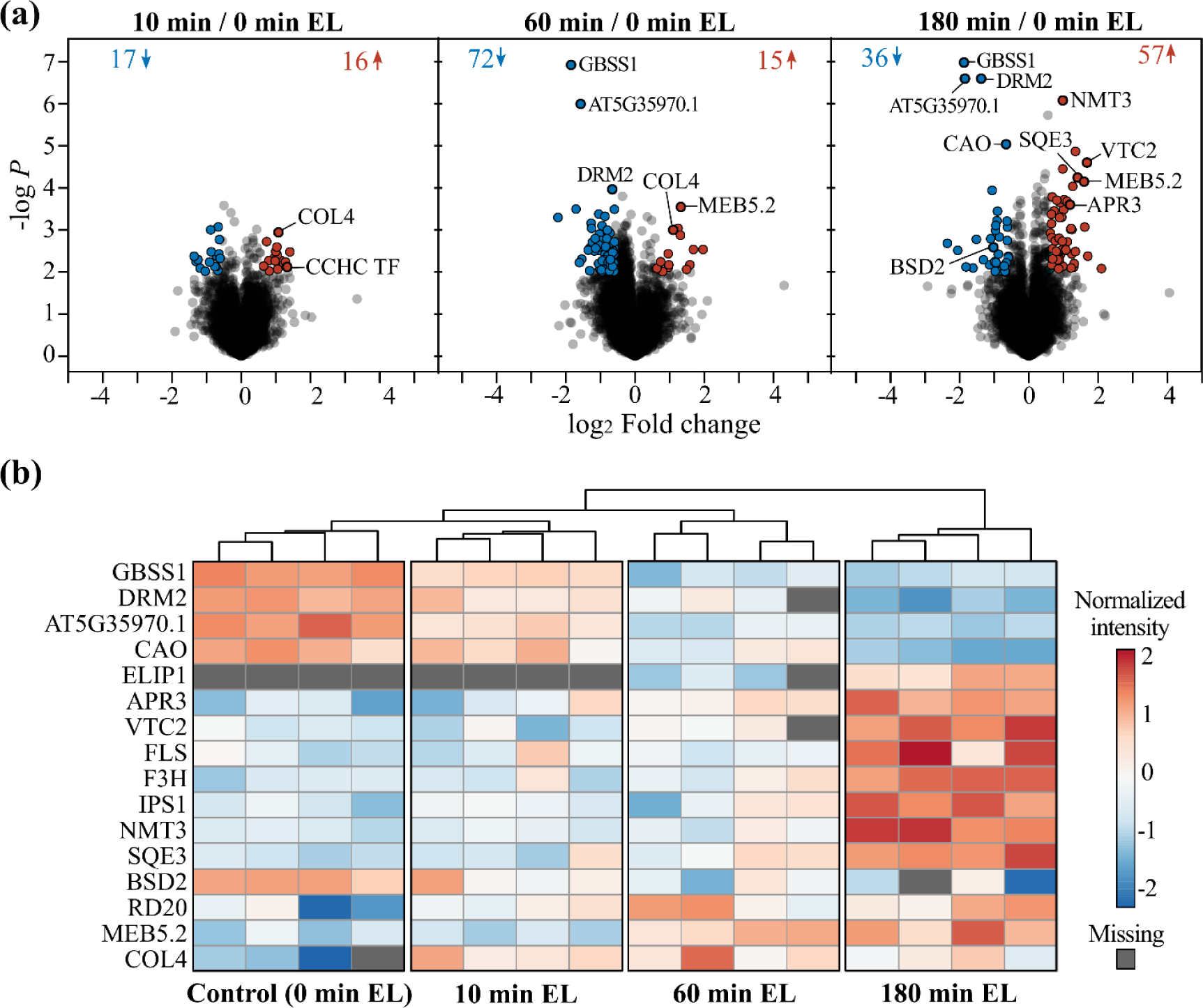
Altered protein abundance under excess light. (**A**) Volcano plots of differential protein expression in EL conditions analyzed with MSstats (*38*). Differentially abundant proteins were selected at a *P* value ≤ 0.01 and an absolute fold change > 1.5, indicated in blue (repressed) and red (induced). (**B**) Protein abundance heatmap created using ClustVis (*78*). Log10 protein abundance after MSstats normalization were centered and unit variance scaling was applied (rows), missing values are colored grey. Samples are clustered using Euclidean distance and average linkage. Abbreviations: APR3, ADENOSINE PHOSPHOSULFATE REDUCTASE 3 (UniProtKB P92980); BSD2, BUNDLE SHEATH DEFECTIVE 2 (Q9SN73); CAO, CHLOROPHYLLIDE A OXYGENASE (Q9MBA1); COL4, CONSTANS-LIKE 4 (Q9LTB8); DRM2, DORMANCY-ASSOCIATED GENE 2 (P93017); ELIP1, EARLY LIGHT-INDUCED PROTEIN 1 (P93735); F3H, FLAVANONE 3-HYDROXYLASE (Q9S818); FLS1, FLAVONOL SYNTHASE 1 (Q96330); GBSS1, GRANULE-BOUND STARCH SYNTHASE 1 (Q9M); IPS1, INOSITOL 3-PHOSPHATE SYNTHASE 1 (P42801); NMT3, PHOSPHOETHANOLAMINE N-METHYLTRANSFERASE 3 (Q9C6B9); RD20, RESPONSIVE TO DESICCATION 20 (O22788); SQE3, SQUALENE OXIDASE 3 (Q8VYH2); VTC2, VITAMIN C DEFECTIVE 2 (Q8RWE8).

Furthermore, the redox enzymes ADENOSINE PHOSPHOSULFATE REDUCTASE 3 (APR3) and VITAMIN C DEFECTIVE 2 (VTC2) were induced after 180 minutes of EL (2.27- and 3.20-fold induced, respectively). APR3 is a chloroplastic enzyme involved in Cys biosynthesis that is strongly induced under H_2_O_2_ stress, likely promoting glutathione and Cys synthesis under these conditions (*44*). VTC2 is a rate-limiting enzyme in ascorbate biosynthesis, whose deficiency causes reduced ascorbate content and photo-oxidative stress under EL (*45*). Taken together, increased levels of both redox enzymes likely increase the rate of Cys and ascorbate metabolism to maintain the redox homeostasis under EL stress. In addition, several enzymes involved in plant secondary metabolism were induced after 180 minutes of EL. These include biosynthetic enzymes of the flavonoid biosynthesis pathway, such as FLAVONOL SYNTHASE 1 [FLS1] and FLAVANONE 3-HYDROXYLASE [F3H], which promote the synthesis of protective metabolites like flavonols and anthocyanins under EL (*46*). Increased abundance was also apparent for enzymes part of other metabolic branches, including INOSITOL 3-PHOSPHATE SYNTHASE 1 (IPS1, 2.03-fold induction, myo-inositol biosynthesis), SQUALENE OXIDASE 3 (SQE3, 2.65-fold induced, steroid biosynthesis), and PHOSPHOETHANOLAMINE N-METHYLTRANSFERASE 3 (NMT3, 1.97-fold induced, phosphatidylcholine biosynthesis).

Lastly, certain proteins with significant changes upon EL stress have been already associated with abiotic stress and redox responses. For instance, the chloroplast protein BUNDLE SHEATH DEFECTIVE 2, suggested to reduce and re-activate oxidized ribulose bisphosphate carboxylase (*47*), was found at lower levels under EL. In this respect, four quantified Cys residues (Cys172, Cys221, Cys427 and Cys259) in Rubisco, were strongly reduced in each sample (< 10% oxidation, Data S1). Protein levels of the transcription factor (TF) CONSTANS-LIKE 4 (COL4), the Ca^2+^-binding protein RESPONSIVE TO DESICCATION 20 (RD20), and the uncharacterized MEB5.2 protein were increased upon EL, in line with their known transcriptional upregulation in response to abiotic stress (*48–50*).

## Discussion

Protein redox regulation is an essential post-translational regulation layer in cellular signaling, with cysteines taking a prominent position due to their intrinsic high reactivity. To comprehensively record protein abundance and the oxidation degree of Cys between and in proteomes, we developed a novel approach termed CysQuant that combines label-free shotgun proteomics with labeled quantification of isotopologous IAM. CysQuant is compatible with a wide range of mass spectrometry platforms for assessing the Cys modification status.

As originally implemented in the OxiCAT method (*15*), our workflow determines a degree of Cys oxidation, which is very valuable information for biological interpretation (*13, 14*). While the majority of Cys thiols were in a reduced state averaging ∼18% of oxidation, on average 82.5% oxidation was evident for cysteines that are part of disulfides in AlphaFold2 predicted structures (*33, 51*), which serves as a biological quality check for our redox proteomics method. Such AlphaFold2-based annotation has been of value to point to potentially regulatory disulfides in proteins lacking crystal structures. One limitation of our workflow is that TCEP reduction used in this study indiscriminately reduces various oxidative modifications, such as *S*-nitrosylation, *S*-sulfenylation, *S*-glutathionylation, and others, using more selective reductive agents such as ascorbate (*S*-nitrosothiols) (*52*), hydroxylamine (*S*-acylation) (*53*), or arsenite (*S*-sulfenylation) (*54*) would also be compatible with the CysQuant workflow. Noteworthy, in case that Cys are subject to sulfinic and sulfonic acid modifications (–SO_2_H and –SO_3_H) that cannot be reversed by TCEP, the calculated oxidation degrees would be an underestimation. However, a label-free search specifying these overoxidation forms identified only a handful of *S*-sulfonylation and ∼100 *S*-sulfinylation sites (Data S4), including active sites of proteins such as GLUTATHIONE PEROXIDASE1 and PEROXIDOXIN Q.

Isotopologous IAM reagents employed in this study have thus far been used to estimate Cys oxidation degrees for purified proteins (*55, 56*) and for labeling reduced Cys across samples to monitor Cys oxidation indirectly in the SICyLIA workflow (*29*). In the latter workflow, oxidation levels were normalized by protein abundance recorded by stable isotope dimethyl labelling. In the CysQuant workflow, protein abundance is quantified in a label-free manner, yet no additional normalization is required as ratios of reduced and reversibly oxidized Cys levels can be estimated directly. In contrast to other isotopologous cysteine mass tags (*15, 16, 57*), a distinct advantage of IAM labeling is that the fragmentation, retention time and other characteristics of IAM-modified peptides can be accurately predicted using available predictors due to its routine incorporation in proteomic workflows. This was capitalized on here, using DIA-NN with *in silico* predicted spectral libraries, providing a convenient and widely accessible DIA-only approach with labeled quantification via its plexDIA module. Moreover, implementing this workflow on MS instruments incorporating ion mobility separation, e.g. DIA-PASEF (*58*), seems a promising avenue to further achieve higher confident quantification with reduced ion interference, improved sensitivity and greater proteome-depth. Importantly, CysQuant does not compromises protein quantification and can thus be regarded as an additional quantitative dimension of Cys oxidation, which can be of interest to a multitude of experimental set-ups including redox physiological processes, but also biomarker discovery.

We used CysQuant to study the excess light response of plant seedlings and identified highly significant reduction of regulatory disulfides in carbon fixation enzymes, which are arguably the best-characterized redox mechanisms in photosynthetic organisms (*36, 59*). In total, four out of six redox-regulated Calvin-Benson cycle enzymes and accessory proteins were successfully recorded by CysQuant (Fig. S8). These included regulatory disulfides of the CP12-2/PRK/GAPB ternary complex and in the C-terminal domain of the RCA-α isoform, which are all reduced and activated by Trxs in a light-dependent manner (*30, 35, 60–62*). Importantly, our analysis discovered reduction of disulfides and potential redox regulation of additional carbon assimilation enzymes in the chloroplast. Most closely connected to the Calvin-Benson cycle is PHOSPHOGLUCOSE ISOMERASE 1 (PGI1), which converts fructose-6-phosphate to glucose-6-phosphate and sequestrates carbon towards other pathways such as starch synthesis. Interestingly, PGI1 and other starch biosynthetic enzymes are inactivated by light (*63*), which thus might be (partially) dependent on the observed light-induced reduction of a putative Cys548-Cys606 disulfide in an extended C-terminal domain of PGI1 (see Fig. S9A). Furthermore, we identified a ∼25% significant reduction of a putative disulfide Cys143-Cys149 in chloroplastic PPDK REGULATORY PROTEIN 1 (RP1) under EL. RP1 regulates the phosphorylation state of a catalytic residue in PYRUVATE, ORTHOPHOSPHATE DIKINASE (PPDK), an enzyme that regenerates phospho*enol*pyruvate in C4 metabolism, in a light-dependent manner suggested to be independent of redox (*64, 65*). The AlphaFold2 predicted disulfide corresponds to the flexible Cys165-Cys171 disulfide that stabilized the dimer formation of the maize orthologue PDRP1 (*66*) (Fig. S9A). Similarly, the C-terminal domain of PGI1 was shown to facilitate dimer formation (*67*) (Fig. S9B). Like PGI1, follow-up biochemical experiments are needed to address the potential redox regulation of RP1 in Arabidopsis (or other plants with C4 metabolism). Next to reduction, several Cys showed increased oxidation upon EL, including an increase in oxidation of TPC1 Cys392 at 10 min EL. TPC1 facilitates systemic Ca^2+^ and ROS signaling waves (*68*), which propagate within minutes, and such rapid Cys could contribute to cross-talk and integration of Ca^2+^ and ROS signals. Nevertheless, CysQuant is a powerful hypothesis formulator of protein redox regulation in physiological conditions, especially in combination with the structural perspective offered by AlphaFold2 predictions.

In contrast to MS1-level quantification, MS2-level quantification can be hindered in CysQuant by the lack of differential fragment ions (containing IAM^0^/IAM^4^). Lysine/arginine labelling strategies such as SILAC or mTRAQ yield C-terminally labelled tryptic peptides with thus differentially labelled y-ion fragment series. In contrast, Cys residues do not typically occur at the C-terminus of tryptic peptides, and the number of quantified differentiating fragment ions can be a restricting factor for MS2-level quantification (examples; see Fig. S10A-B). For instance, only 228 peptide precursor ions presented two differentiating fragment ions (out of the six highest abundant) in three out of four replicates in the control condition – which is drastically lower than the 1460 MS1-level quantified ratios (not including L/H-only channel precursors). Nevertheless, in some cases, the presence of differentiating fragment ions can inform on specific Cys within isobaric peptides containing multiple Cys. DIA analysis has been applied before in such a context for histone modifications (*69*) or phosphorylation site assignment (*70*), and can also be achieved with CysQuant MS2 level quantification in some cases. For instance, FTSH5 INTERACTING PROTEIN contains a ‘CPEC’ motif of a zinc-finger domain for which the second Cys is quantified by consecutive y-ions y8+, y9+, y10+ and y10++ (Fig. S10C). While MS1-level plexDIA quantification is recommended for proteome-wide analysis when using isotopologous IAM labels, MS2-level quantification can still be of value in certain cases.

In summary, we present here a novel redox proteome profiling method, CysQuant, that is easy adoptable for DDA and/or DIA analysis and that simultaneously profiles protein abundance and Cys oxidation degrees. By using CysQuant to study the excess light response in Arabidopsis seedlings, we detected unbiasedly well-established redox regulation in the chloroplast and discovering yet uncharacterized redox regulation for other chloroplast enzymes. This method and its findings serves as a valuable scientific resource in the field redox biology. We believe CysQuant method to be instrumental for diverse Cys proteome studies, such as Cys oxidation profiling in entire proteomes or recombinant proteins, profiling substrates of thioredoxin or other redox-active proteins, or Cys ligand screening and reactivity profiling.

## Materials and Methods

### Plant growth conditions and stress treatments

Arabidopsis (*Arabidopsis thaliana* L. Heynh.) cv. Columbia-0 (Col-0) was grown in a controlled-environment growth chamber (Weiss Technik; 16 h [50 μmol m^−2^ s^−1^] light/8 h dark, 21 °C, 60–65% humidity). For excess light (EL) treatment, trays of 23-day-old plants were shifted within the same growth chamber to an upper shelf at a 1,000 μmol m^−2^ s^−1^ irradiance. Changes in photosystem II maximum efficiency (F_v_’/F_m_’) signals were quantified by an Imaging PAMM-series (MAXI version) chlorophyll fluorometer and the ImagingWin software application (Heinz Walz).

### CysQuant differential thiol labeling

Rosettes of 23-day-old seedlings grown on soil were collected and frozen in liquid nitrogen. After the plant material had been ground to a fine powder with mortar and pestle with liquid nitrogen, the frozen tissue was suspended in 20% (v/v) trichloroacetic acid (TCA) and vortexed vigorously. Samples were lysed and homogenized by sonication and left on ice for 30 min. Samples were subsequently centrifuged at 15,000 rpm for 30 min at 4 °C. The pellet was washed once with 10% (v/v) TCA and two times with cold (−20 °C) acetone. After centrifugation and air evaporation to remove residual acetone, the precipitate was resuspended in protein extraction buffer containing 10% (w/v) SDS, 5 mM EDTA, and 40 mM (light) IAM (Sigma-Aldrich) in 100 mM HEPES, pH 7.5. Samples were vortexed vigorously and incubated for 1 h at 37 °C in the dark. Protein concentrations were measured using the BCA protein assay kit (Thermo Scientific). Further protein purification was performed using an adapted version of the S-Trap^TM^ mini column workflow (ProtiFi). Samples were acidified using 12% phosphoric acid (diluted in water), vortexed and mixed with six times the sample volume of binding buffer (100 mM HEPES buffer containing 90% methanol, pH 7.5). For each sample, a total of 300 mg of protein was loaded on the S-Trap column and centrifuged at 10,000*g* for 30 s to trap proteins. Trapped proteins were cleaned three times with 400 µL binding buffer. Afterwards, oxidized thiols were reduced on-column by adding 500 mM tris(2-carboxyethyl)phosphine (TCEP) (Sigma-Aldrich) prepared freshly in a 100 mM HEPES buffer (pH 7.5) and incubating the S-Trap column for 1 h at 37 °C. The reaction was stopped by centrifugation at 10,000*g* for 30 s. Reduced thiols were then labelled with heavy IAM^4^ (^13^C_2_D_2_H_2_INO-IAM, Sigma-Aldrich cat. no. 721328) by adding 40 mM IAM^4^ in 100 mM HEPES (pH 7.5) on the S-Trap columns and incubating the column for 1 h at 37 °C in the dark. Protein samples were cleaned by adding 50 mM NH_4_HCO_3_ buffer followed with centrifugation at 10,000*g* for 30 s (repeated three times) to remove residual TCEP and IAM^4^. For each protein sample, on-column trypsin digestion was performed by adding by mass spectrometry grade trypsin/Lys-C mix (Promega) in a 50 mM NH_4_HCO_3_ buffer overnight at 37 °C.

### LC-MS/MS analysis

From each sample, 0.5 µL was injected onto the column for LC-MS/MS analysis using an Ultimate 3000 RSLC nano system in-line connected to a Q Exactive HF BioPharma mass spectrometer (Thermo Fisher Scientific). Trapping was performed at 20 μL/min for 2 min in loading solvent A (0.1% TFA in water/ACN (99.5:0.5, [v/v]) on a 5 mm trapping column (Thermo Fisher Scientific, 300 μm internal diameter). After flushing from the trapping column, the peptides were loaded and separated on an analytical 250 mm Aurora Ultimate, 1.7 µm C18, 75 µm inner diameter (Ionopticks) kept at a constant temperature of 45 °C. Peptides were eluted using a non-linear gradient starting at 0.5% MS solvent B reaching 26% MS solvent B (0.1% FA in acetonitrile) in 82.5 min, 44% MS solvent B in 96.5 min, 56% MS solvent B in 100 minutes followed by a 5-minutes wash at 56% MS solvent B and re-equilibration with MS solvent A (0.1% FA in water). For both analyses, a pneu-Nimbus dual column ionization source was used (Phoenix S&T), at a spray voltage of 2.3 kV and a capillary temperature of 275 °C.

#### Data-dependent acquisition

The mass spectrometer was operated in data-dependent mode, automatically switching between MS and MS/MS acquisition for the 12 most abundant ion peaks per MS spectrum. Full-scan MS spectra (350-1500 *m/z*) were acquired at a resolution of 120,000 in the Orbitrap analyzer after accumulation to a target value of 3,000,000 with a maximum ion time of 60 ms. The 12 most intense ions above a threshold value of 13,000 were isolated with a width of 1.5 *m/z* for fragmentation at a normalized collision energy of 28% after filling the trap at a target value of 100,000 for a maximum of 80 ms. MS/MS spectra (200-2000 *m/z*) were acquired at a resolution of 15,000 in the Orbitrap analyzer. QCloud (*71, 72*) was used to control instrument longitudinal performance during the project.

#### Data-independent acquisition

The mass spectrometer was operated in data-independent mode, automatically switching between MS and MS/MS acquisition. Full-scan MS spectra (375-1500 *m/z*) were acquired at a precursor resolution of 60,000 at 200 *m/z* in the Orbitrap analyser after accumulation to a target value of 5E6 with a maximum ion time of 50 ms. After each MS1 acquisition, thirty 10-*m/z* isolation windows were sequentially selected by the quadrupole for HCD fragmentation at a NCE of 30% after filling the trap at a target value of 3E6 for a maximum injection time of 45 ms. MS2 spectra were acquired at a resolution of 15,000 at 200 *m/z* in the Orbitrap analyser without multiplexing. The isolation windows were acquired over a mass range of 400-900 *m/z*. The isolation windows were created with the Skyline software tool (v3.6).

### DDA data analysis

Raw data files were searched with MaxQuant (version 2.3.1.0) (*27*) against the Arabidopsis UniProtKB reference proteome (UP000006548, 27,474 proteins) and MaxQuant built-in contaminant proteins. We used the enzymatic rule of Trypsin/P with a maximum of two missed cleavages. To quantify H/L ratios of IAM-modified peptides, light (+57.021 Da, IAM^0^) and heavy IAM (+61.041 Da, IAM^4^) were set as labels similar as described (*29*). The re-quantify option was enabled to rescue incomplete isotope pattern pairs (*28*). In parallel, a label-free MaxQuant search was performed with IAM^0^ and IAM^4^ specified as variable modifications. For both searches, N-terminal protein acetylation and methionine oxidation were set as variable modifications. To augment peptide quantification, we enabled matching-between-runs (default match time window of 0.7 min, alignment time window of 20 min), the second peptide search, and the LFQ algorithm using default settings. All FDR thresholds were kept at the default threshold of 1%. For other search parameters not specified here, default MaxQuant settings were used. For identification of overoxidized Cys, S-sulfinylation (+31.9898 Da) and S-sulfonylation (+47.9847 Da) were specified as additional variable modifications.

### DIA data analysis

#### Spectral library generation

In silico-predicted spectral libraries constructed by DIA-NN (version 1.8.2 beta 22) (*21*, *25, 73*) based on the UniProtKB reference proteome FASTA file (see DDA analysis) were used. Peptide length was set at 7 to 30 residues, with a precursor *m/z* ranging between 395-905 *m/z*. N-terminal methionine excision and a single missed cleavage by trypsin were allowed. In total, 2,440,135 precursors for 27,416 proteins were generated.

#### plexDIA analysis

RAW files were processed by the plexDIA module of DIA-NN (version 1.8.2 beta 22) (*21*,*25,73*) for protein abundance and Cys oxidation degree profiling using the *in silico* predicted spectral library described above. Instead of lysine/arginine chemical mass tags, isotopologous IAM modifications were here specified as chemical mass tags using the parameters ‘*–fixed-mod “IAM,0.0,C,label” –lib-fixed-mod IAM –channels “IAM,L,C,57.021464; IAM,H,C,61.04073”*’. Other parameters were identical as described before (*21*). The MS1 mass accuracy was set at 7 ppm and scan window at 6 as recommended from initial pilot searches. Matching-between-runs was enabled and the heuristic protein interference was set off.

### Cysteine oxidation degree

The degree of Cys oxidation was calculated using the following formula:

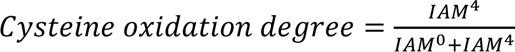

with IAM^0^ and IAM^4^ representing the reduced (light, L) and oxidized (heavy, H) MS1 intensities. In case of MaxQuant DDA analysis, the (non-normalized) H/L ratios for peptides containing one Cys were used from the ‘peptides.txt’ output of the labelled search. Note that we did not retain light (L) or heavy (H) channel only quantifications, as this resulted in a dramatic increase in variation. For the DIA analysis, the main DIA-NN output report was used to estimate the Cys oxidation degree at MS1 and MS2 level. For peptide precursors with quantified MS1 areas (Ms1.area > 0) and a Ms1 profile correlation ≥ 0.2 in the light and heavy channel, oxidation degrees were retained if either the L or H precursor had a channel and translated Q value ≤ 1% (Data S1). For MS2 level quantification, we summed up the L/H areas of the six most abundant ion fragments if containing a Cys (i.e. differentiating fragment ion) and having a fragment correlation ≥ 0.5 for both the L and H precursor. MS2-level oxidation degrees were only reported for oxidation degrees for peptide precursors with at least two differential fragment ions and if either the L or H precursor had a channel and translated Q value ≤ 1%.

Unlike multiplexed protein abundance, Cys redox labeling often presents extreme ratios due to the presence of strongly reduced or strongly oxidized thiols (e.g. in structural disulfides). To retain confident cases of precursors solely present in the L channel (i.e. fully reduced) or H channel (i.e. fully oxidized) in the MS1-level analysis, we required an additional Ms1 and Ms2 channel evidence filter ≥ 0.5 and a Ms1 intensity surpassing the 25% quantile of the run intensities recorded for Cys peptides with L/H ratios. This intensity-based filter was used to avoid the missing L or H channel to fall under the detection limit. Note that this 25% quantile intensity threshold (nor less strict thresholds) only retained twelve L or H channel only quantifications in the MaxQuant ‘peptides.txt’.

### Differential protein abundance

For uniform analysis of differential protein abundance in DDA and DIA analysis, we used the MSstats package (version 4.7.1) (*38*). From the MaxQuant label-free search, with IAM^0^ and IAM^4^ as variable modifications, the ‘proteinGroups.txt’ and ‘evidence.txt’ file were converted to a MSstats input report using the MSstatsConvert package (version 1.4.1). We discarded peptide precursors containing Cys (via the ‘pattern_filtering’ parameter in the *MSstatsPreprocess* function) or assigned ambiguously to multiple proteins. MS2 fragment quantifications (‘Fragment.Quant.Raw’) of non-Cys peptide precursors with a global Q-value ≤ 1% of the DIA-NN report were used to construct a MSstats input report. For both DDA and DIA inputs, we opted for data processing retaining high quality features (*74*) via adapting the parameters: (1) ‘*–featureSubset = highQuality*’ (2) ‘*–remove_uninformative_feature_outlier = TRUE’*, (3) ‘*–remove50missing = TRUE’*, and (4) ‘*–min_feature_count = 2’*. Afterwards, pairwise comparisons of interest, i.e. excess light treatment versus respective control conditions, were performed using the recommended guidelines. Thresholds for selecting significant differential proteins were *P* ≤ .01 and absolute fold change ≥ 1.5.

### Statistical analysis

Quantitative data of cysteine oxidation degrees are presented as histogram, boxplot or violin plot, with sample sizes indicated in the figure. Differential analysis of protein abundance was performed using MSstats (*38*) as described above. To assess changes in the degree of Cys oxidation, pairwise comparisons were made between each EL condition and the control condition using a two sample Welch’s t-test in R (version 4.3.1). Absolute Cys oxidation degree changes greater than 5% with a *P* value ≤ 0.05 were considered as significant.

## Acknowledgments

We are grateful to Robin Pottie for helping with the chlorophyll fluorescent measurements.

## Author contributions

Conceptualization: PW, KG, AS, JH. Methodology: JH, PW, AS, VD, KG. Investigation: JH, AS, PW. Visualization: PW. Resources: KG, VD, FI, FVB. Supervision: KG. Writing—original draft: PW, JH. Writing—review & editing: JH, AS, VD, FI, FVB, KG, PW.

## Data and materials availability

The DIA-NN 1.8.2 beta 22 algorithm (*73*) is available at the OSF repository https://osf.io/q8kfc/?view_only=5e77d3c62563468280fd09265583dbbd. The raw proteomics data sets, spectral library, FASTA, and MaxQuant/plexDIA results are deposited to the ProteomeXchange Consortium via the PRIDE (*79*) partner repository with the dataset identifier PXD043895. The Python script used to calculate cysteine oxidation degrees from the plexDIA analysis is available at https://github.com/patrick-willems/CysQuant. All data are available in the main text or the supplementary materials.

## References

1. P. Willems, F. Van Breusegem, J. Huang, Contemporary proteomic strategies for cysteine redoxome profiling. Plant Physiol 186, 110–124 (2021).

2. A. J. Meyer, J. Riemer, N. Rouhier, Oxidative protein folding: state-of-the-art and current avenues of research in plants. New Phytol 221, 1230–1246 (2019).

3. G. Bi et al., The cytosolic thiol peroxidase PRXIIB is an intracellular sensor for H_2_O_2_ that regulates plant immunity through a redox relay. Nat Plants 8, 1160–1175 (2022).

4. E. A. Veal et al., A 2-Cys peroxiredoxin regulates peroxide-induced oxidation and activation of a stress-activated MAP kinase. Mol Cell 15, 129–139 (2004).

5. A. Delaunay, D. Pflieger, M. B. Barrault, J. Vinh, M. B. Toledano, A thiol peroxidase is an H2O2 receptor and redox-transducer in gene activation. Cell 111, 471–481 (2002).

6. Y. Meyer, C. Belin, V. Delorme-Hinoux, J. P. Reichheld, C. Riondet, Thioredoxin and glutaredoxin systems in plants: molecular mechanisms, crosstalks, and functional significance. Antioxid Redox Signal 17, 1124–1160 (2012).

7. C. H. Foyer, G. Noctor, Ascorbate and glutathione: the heart of the redox hub. Plant Physiol 155, 2–18 (2011).

8. J. Huang et al., Mining for protein S-sulfenylation in Arabidopsis uncovers redox-sensitive sites. Proc Natl Acad Sci U S A 116, 21256–21261 (2019).

9. S. Akter et al., Chemical proteomics reveals new targets of cysteine sulfinic acid reductase. Nat Chem Biol 14, 995–1004 (2018).

10. U. Seneviratne et al., S-nitrosation of proteins relevant to Alzheimer’s disease during early stages of neurodegeneration. Proc Natl Acad Sci U S A 113, 4152–4157 (2016).

11. B. Wei et al., Identification of Sulfenylated Cysteines in Arabidopsis thaliana Proteins Using a Disulfide-Linked Peptide Reporter. Front Plant Sci 11, 777 (2020).

12. I. S. Arts, D. Vertommen, F. Baldin, G. Laloux, J. F. Collet, Comprehensively Characterizing the Thioredoxin Interactome In Vivo Highlights the Central Role Played by This Ubiquitous Oxidoreductase in Redox Control. Mol Cell Proteomics 15, 2125–2140 (2016).

13. G. Prus, A. Hoegl, B. T. Weinert, C. Choudhary, Analysis and Interpretation of Protein Post-Translational Modification Site Stoichiometry. Trends Biochem Sci 44, 943–960 (2019).

14. J. Duan, M. J. Gaffrey, W. J. Qian, Quantitative proteomic characterization of redox-dependent post-translational modifications on protein cysteines. Mol Biosyst 13, 816–829 (2017).

15. L. I. Leichert et al., Quantifying changes in the thiol redox proteome upon oxidative stress in vivo. Proc Natl Acad Sci U S A 105, 8197–8202 (2008).

16. S. Doron et al., SPEAR: A proteomics approach for simultaneous protein expression and redox analysis. Free Radic Biol Med 176, 366–377 (2021).

17. H. Xiao et al., A Quantitative Tissue-Specific Landscape of Protein Redox Regulation during Aging. Cell 180, 968–983 e924 (2020).

18. T. Nietzel et al., Redox-mediated kick-start of mitochondrial energy metabolism drives resource-efficient seed germination. Proc Natl Acad Sci U S A 117, 741–751 (2020).

19. C. B. Messner et al., Ultra-High-Throughput Clinical Proteomics Reveals Classifiers of COVID-19 Infection. Cell Syst 11, 11–24 e14 (2020).

20. Extending the sensitivity, consistency and depth of single-cell proteomics. Nat Methods 20, 649-650 (2023).

21. J. Derks et al., Increasing the throughput of sensitive proteomics by plexDIA. Nat Biotechnol 41, 50–59 (2023).

22. K. Barkovits et al., Reproducibility, Specificity and Accuracy of Relative Quantification Using Spectral Library-based Data-independent Acquisition. Mol Cell Proteomics 19, 181–197 (2020).

23. S. I. Anjo et al., oxSWATH: An integrative method for a comprehensive redox-centered analysis combined with a generic differential proteomics screening. Redox Biol 22, 101130 (2019).

24. F. Yang, G. Jia, J. Guo, Y. Liu, C. Wang, Quantitative Chemoproteomic Profiling with Data-Independent Acquisition-Based Mass Spectrometry. J Am Chem Soc 144, 901–911 (2022).

25. V. Demichev, C. B. Messner, S. I. Vernardis, K. S. Lilley, M. Ralser, DIA-NN: neural networks and interference correction enable deep proteome coverage in high throughput. Nat Methods 17, 41–44 (2020).

26. M. HaileMariam et al., S-Trap, an Ultrafast Sample-Preparation Approach for Shotgun Proteomics. J Proteome Res 17, 2917–2924 (2018).

27. J. Cox, M. Mann, MaxQuant enables high peptide identification rates, individualized p.p.b.-range mass accuracies and proteome-wide protein quantification. Nat Biotechnol 26, 1367–1372 (2008).

28. S. Tyanova, M. Mann, J. Cox, MaxQuant for in-depth analysis of large SILAC datasets. Methods Mol Biol 1188, 351–364 (2014).

29. J. van der Reest, S. Lilla, L. Zheng, S. Zanivan, E. Gottlieb, Proteome-wide analysis of cysteine oxidation reveals metabolic sensitivity to redox stress. Nat Commun 9, 1581 (2018).

30. K. Yoshida, Y. Yokochi, K. Tanaka, T. Hisabori, The ferredoxin/thioredoxin pathway constitutes an indispensable redox-signaling cascade for light-dependent reduction of chloroplast stromal proteins. J Biol Chem 298, 102650 (2022).

31. P. Schurmann, B. B. Buchanan, The ferredoxin/thioredoxin system of oxygenic photosynthesis. Antioxid Redox Signal 10, 1235–1274 (2008).

32. N. R. Baker, Chlorophyll fluorescence: a probe of photosynthesis in vivo. Annu Rev Plant Biol 59, 89–113 (2008).

33. P. Willems, J. Huang, J. Messens, F. Van Breusegem, Functionally annotating cysteine disulfides and metal binding sites in the plant kingdom using AlphaFold2 predicted structures. Free Radic Biol Med 194, 220–229 (2023).

34. Z. J. Wehrspan, R. T. McDonnell, A. H. Elcock, Identification of Iron-Sulfur (Fe-S) Cluster and Zinc (Zn) Binding Sites Within Proteomes Predicted by DeepMind’s AlphaFold2 Program Dramatically Expands the Metalloproteome. J Mol Biol 434, 167377 (2022).

35. N. Zhang, A. R. Portis, Jr., Mechanism of light regulation of Rubisco: a specific role for the larger Rubisco activase isoform involving reductive activation by thioredoxin-f. Proc Natl Acad Sci U S A 96, 9438–9443 (1999).

36. L. Michelet et al., Redox regulation of the Calvin-Benson cycle: something old, something new. Front Plant Sci 4, 470 (2013).

37. D. Wang, A. R. Portis, Jr., A novel nucleus-encoded chloroplast protein, PIFI, is involved in NAD(P)H dehydrogenase complex-mediated chlororespiratory electron transport in Arabidopsis. Plant Physiol 144, 1742–1752 (2007).

38. M. Choi et al., MSstats: an R package for statistical analysis of quantitative mass spectrometry-based proteomic experiments. Bioinformatics 30, 2524–2526 (2014).

39. K. Denyer, D. Waite, A. Edwards, C. Martin, A. M. Smith, Interaction with amylopectin influences the ability of granule-bound starch synthase I to elongate malto-oligosaccharides. Biochem J 342 Pt 3, 647–653 (1999).

40. A. L. Harper et al., Chlorophyllide a Oxygenase mRNA and Protein Levels Correlate with the Chlorophyll a/b Ratio in Arabidopsis thaliana. Photosynth Res 79, 149–159 (2004).

41. C. Hutin et al., Early light-induced proteins protect Arabidopsis from photooxidative stress. Proc Natl Acad Sci U S A 100, 4921–4926 (2003).

42. S. Rossini et al., Suppression of both ELIP1 and ELIP2 in Arabidopsis does not affect tolerance to photoinhibition and photooxidative stress. Plant Physiol 141, 1264–1273 (2006).

43. T. Tzvetkova-Chevolleau et al., The light stress-induced protein ELIP2 is a regulator of chlorophyll synthesis in Arabidopsis thaliana. Plant J 50, 795–809 (2007).

44. G. Queval et al., H2O2-activated up-regulation of glutathione in Arabidopsis involves induction of genes encoding enzymes involved in cysteine synthesis in the chloroplast. Mol Plant 2, 344–356 (2009).

45. P. Muller-Moule, T. Golan, K. K. Niyogi, Ascorbate-deficient mutants of Arabidopsis grow in high light despite chronic photooxidative stress. Plant Physiol 134, 1163–1172 (2004).

46. G. E. Araguirang, A. S. Richter, Activation of anthocyanin biosynthesis in high light - what is the initial signal? New Phytol 236, 2037–2043 (2022).

47. J. Tominaga, S. Takahashi, A. Sakamoto, H. Shimada, Arabidopsis BSD2 reveals a novel redox regulation of Rubisco physiology in vivo. Plant Signal Behav 15, 1740873 (2020).

48. J. H. Min, J. S. Chung, K. H. Lee, C. S. Kim, The CONSTANS-like 4 transcription factor, AtCOL4, positively regulates abiotic stress tolerance through an abscisic acid-dependent manner in Arabidopsis. J Integr Plant Biol 57, 313–324 (2015).

49. I. Kalbina, A. Strid, The role of NADPH oxidase and MAP kinase phosphatase in UV-B-dependent gene expression in Arabidopsis. Plant Cell Environ 29, 1783–1793 (2006).

50. S. Takahashi, T. Katagiri, K. Yamaguchi-Shinozaki, K. Shinozaki, An Arabidopsis gene encoding a Ca2+-binding protein is induced by abscisic acid during dehydration. Plant Cell Physiol 41, 898–903 (2000).

51. J. Jumper et al., Highly accurate protein structure prediction with AlphaFold. Nature 596, 583–589 (2021).

52. G. Qin et al., FAT-switch-based quantitative S-nitrosoproteomics reveals a key role of GSNOR1 in regulating ER functions. Nat Commun 14, 3268 (2023).

53. M. Kumar, P. Carr, S. R. Turner, An atlas of Arabidopsis protein S-acylation reveals its widespread role in plant cell organization and function. Nat Plants 8, 670–681 (2022).

54. L. Yu et al., Proteome-wide identification of S-sulfenylated cysteines reveals metabolic response to freezing stress after cold acclimation in Brassica napus. Front Plant Sci 13, 1014295 (2022).

55. Z. Wang et al., Functionality of Redox-Active Cysteines Is Required for Restriction of Retroviral Replication by SAMHD1. Cell Rep 24, 815–823 (2018).

56. N. Burger et al., ND3 Cys39 in complex I is exposed during mitochondrial respiration. Cell Chem Biol 29, 636–649 e614 (2022).

57. M. E. Albertolle, S. M. Glass, E. Trefts, F. P. Guengerich, Isotopic tagging of oxidized and reduced cysteines (iTORC) for detecting and quantifying sulfenic acids, disulfides, and free thiols in cells. J Biol Chem 294, 6522–6530 (2019).

58. F. Meier et al., diaPASEF: parallel accumulation-serial fragmentation combined with data-independent acquisition. Nat Methods 17, 1229–1236 (2020).

59. C. H. Foyer, G. Noctor, Redox homeostasis and antioxidant signaling: a metabolic interface between stress perception and physiological responses. Plant Cell 17, 1866–1875 (2005).

60. L. Marri et al., Prompt and easy activation by specific thioredoxins of calvin cycle enzymes of Arabidopsis thaliana associated in the GAPDH/CP12/PRK supramolecular complex. Mol Plant 2, 259–269 (2009).

61. A. Yu et al., Photosynthetic Phosphoribulokinase Structures: Enzymatic Mechanisms and the Redox Regulation of the Calvin-Benson-Bassham Cycle. Plant Cell 32, 1556–1573 (2020).

62. N. Zhang, P. Schurmann, A. R. Portis, Jr., Characterization of the regulatory function of the 46-kDa isoform of Rubisco activase from Arabidopsis. Photosynth Res 68, 29–37 (2001).

63. B. Heuer, M. J. Hansen, L. E. Anderson, Light modulation of phosphofructokinase in pea leaf chloroplasts. Plant Physiol 69, 1404–1406 (1982).

64. C. J. Chastain et al., The pyruvate, orthophosphate dikinase regulatory proteins of Arabidopsis possess a novel, unprecedented Ser/Thr protein kinase primary structure. Plant J 53, 854–863 (2008).

65. C. J. Chastain, R. Chollet, Regulation of pyruvate, orthophosphate dikinase by ADP-/Pi-dependent reversible phosphorylation in C3 and C4 plants. Plant Physiology and Biochemistry 41, 523–532 (2003).

66. L. Jiang et al., Structural Basis of Reversible Phosphorylation by Maize Pyruvate Orthophosphate Dikinase Regulatory Protein. Plant Physiol 170, 732–741 (2016).

67. F. Gao et al., Engineering of the cytosolic form of phosphoglucose isomerase into chloroplasts improves plant photosynthesis and biomass. New Phytol 231, 315–325 (2021).

68. M. J. Evans, W. G. Choi, S. Gilroy, R. J. Morris, A ROS-Assisted Calcium Wave Dependent on the AtRBOHD NADPH Oxidase and TPC1 Cation Channel Propagates the Systemic Response to Salt Stress. Plant Physiol 171, 1771–1784 (2016).

69. K. A. Krautkramer, L. Reiter, J. M. Denu, J. A. Dowell, Quantification of SAHA-Dependent Changes in Histone Modifications Using Data-Independent Acquisition Mass Spectrometry. J Proteome Res 14, 3252–3262 (2015).

70. S. Sidoli, R. Fujiwara, K. Kulej, B. A. Garcia, Differential quantification of isobaric phosphopeptides using data-independent acquisition mass spectrometry. Mol Biosyst 12, 2385–2388 (2016).

71. C. Chiva et al., QCloud: A cloud-based quality control system for mass spectrometry-based proteomics laboratories. PLoS One 13, e0189209 (2018).

72. R. Olivella et al., QCloud2: An Improved Cloud-based Quality-Control System for Mass-Spectrometry-based Proteomics Laboratories. J Proteome Res 20, 2010–2013 (2021).

73. K. Franziska, L. G. Justus, R. S. Ludwig, D. Vadim, QuantUMS: uncertainty minimisation enables confident quantification in proteomics. bioRxiv, 2023.2006.2020.545604 (2023).

74. T. H. Tsai et al., Selection of Features with Consistent Profiles Improves Relative Protein Quantification in Mass Spectrometry Experiments. Mol Cell Proteomics 19, 944–959 (2020).

75. C. M. Hooper, I. R. Castleden, S. K. Tanz, N. Aryamanesh, A. H. Millar, SUBA4: the interactive data analysis centre for Arabidopsis subcellular protein locations. Nucleic Acids Res 45, D1064–D1074 (2017).

76. C. M. Hooper et al., SUBAcon: a consensus algorithm for unifying the subcellular localization data of the Arabidopsis proteome. Bioinformatics 30, 3356–3364 (2014).

77. E. F. Pettersen et al., UCSF ChimeraX: Structure visualization for researchers, educators, and developers. Protein Sci 30, 70–82 (2021).

78. T. Metsalu, J. Vilo, ClustVis: a web tool for visualizing clustering of multivariate data using Principal Component Analysis and heatmap. Nucleic Acids Res 43, W566–570 (2015).

79. Y. Perez-Riverol et al., The PRIDE database resources in 2022: a hub for mass spectrometry-based proteomics evidences. Nucleic Acids Res 50, D543–D552 (2022).

