## Supplemental Figures S1-S10 for "CysQuant: simultaneous quantification of cysteine oxidation and protein abundance using data dependent or independent acquisition mass spectrometry"

### Supplementary Figures

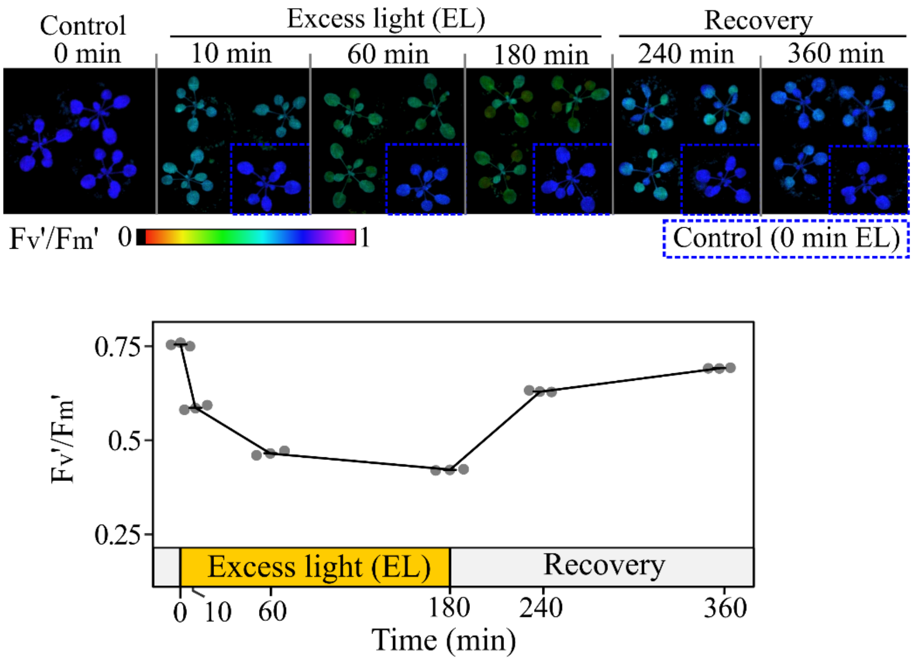


**Fig. S1. Experimental set-up excess light treatment *Arabidopsis thaliana*.**

Twenty-three days old plants were transferred from low light growth conditions to excess light (EL, 50 to 1000 µE $m^{-2}$ $s^{-1}$), displaying a decrease in maximum photosystem II quantum efficiency ($F_v$'/$F_m$') (*N*=3). Shifting back to low light growth conditions recovers $F_v$'/$F_m$'. Control plants not exposed to EL are shown in blue dotted insets. For CysQuant analysis, above-ground seedling tissue was collected at 0, 10, 60 or 180 minutes of EL.

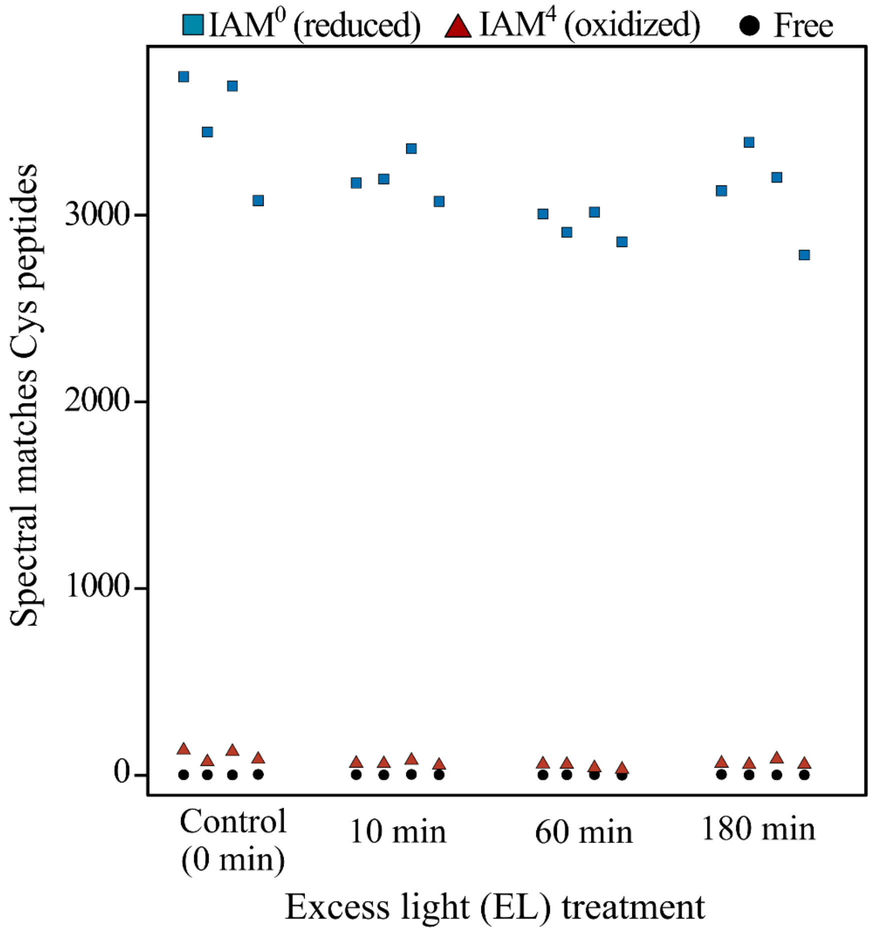


**Fig. S2. Cysteine iodoacetamide labelling efficiency.**

Numbers of spectra matching Cys-containing peptides fully labelled with $IAM^0$ (blue), $IAM^4$ (red) or containing at least one unlabeled Cys in the peptide (black) are shown. Spectral matches were extracted from the ‘msms.txt’ output of the MaxQuant DDA label-free search with $IAM^0$ and $IAM^4$ set as variable modifications.

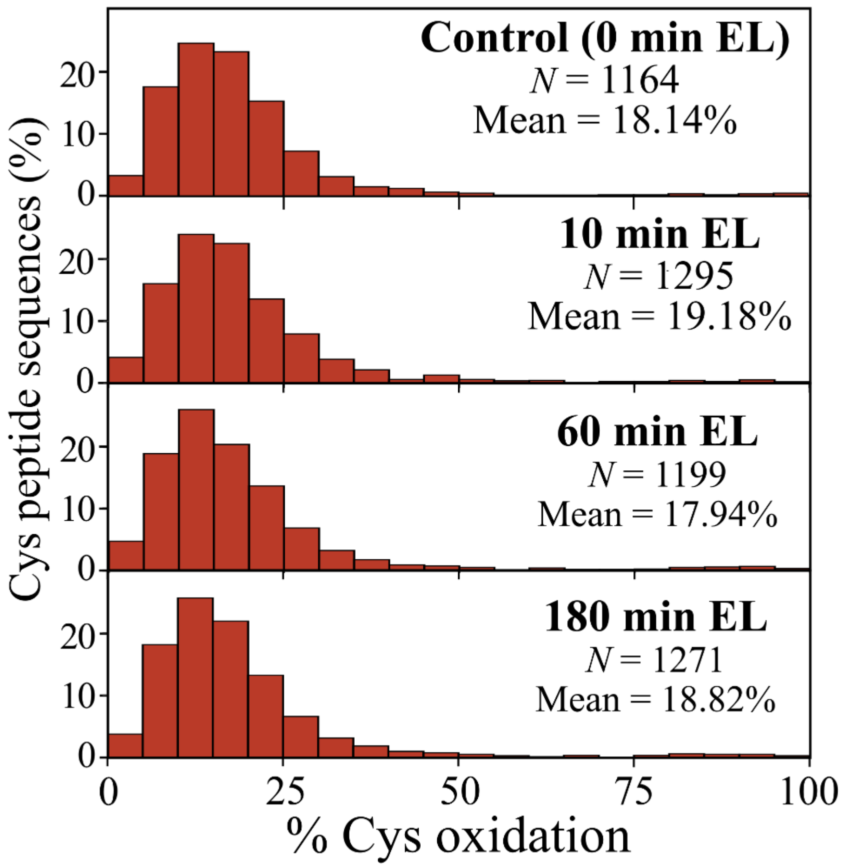


**Fig. S3. Distribution of oxidation degree (%) for cysteines quantified by MaxQuant (DDA) analysis in at least three out of four replicates per condition.**

For detailed overview of quantified cysteines by MaxQuant see Data S1B.

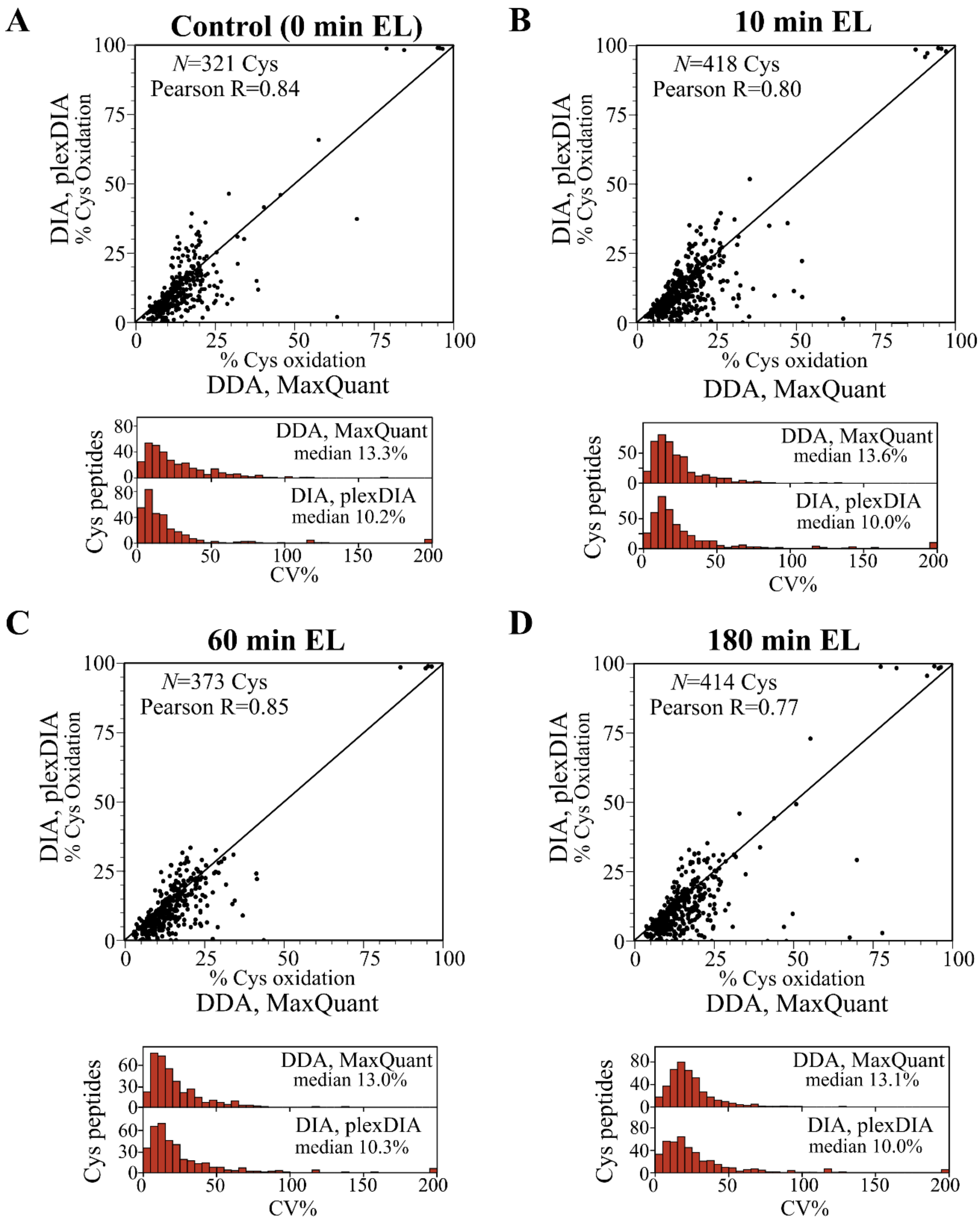


**Fig. S4. Correlation of Cys oxidation degrees measured by MaxQuant (DDA) and plexDIA (DIA) analyses.**

Correlation of the oxidation degrees (%) of Cys quantified by plexDIA and MaxQuant in all four replicates of the control condition (0 minutes of excess light [EL]) (**A**), 10 minutes of EL (**B**), 60 minutes of EL (**C**), and 180 minutes of EL (**D**). Histogram of the coefficient of variation (CV%) for Cys plotted in the correlation analysis (*bottom plots*).

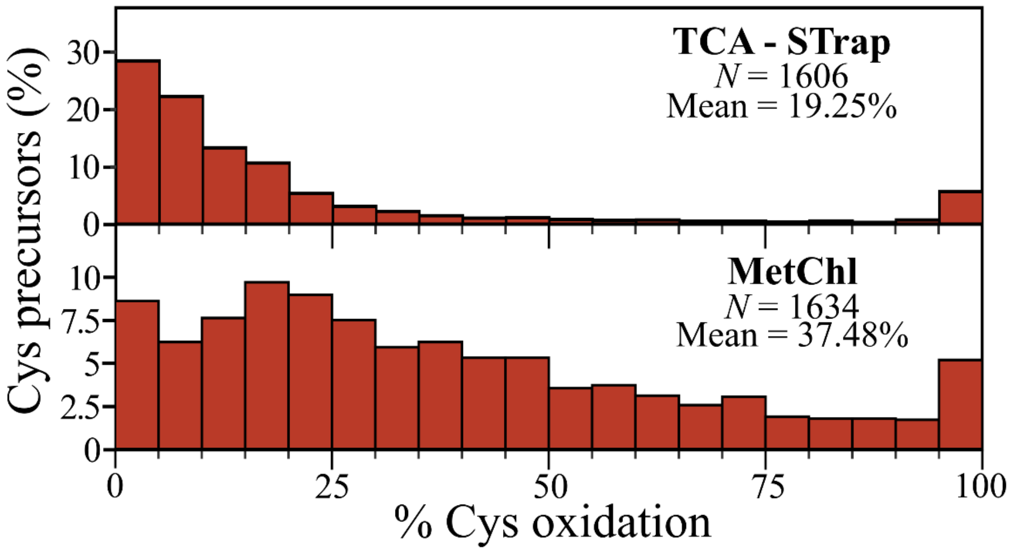


**Fig. S5. Influence of acidic protein extraction on Cys oxidation degrees measured by plexDIA (DIA) analysis.**

Distribution of oxidation degrees (%) for Cys in non-stressed seedlings quantified by plexDIA analysis in at least three out of four replicates after crushing of samples in a trichloroacetic acid (TCA) solution followed by suspension trapping (TCA-STrap) based protein extraction (see Figure 1) or a (non-acidic) methanol-chloroform (MetChl) extraction.

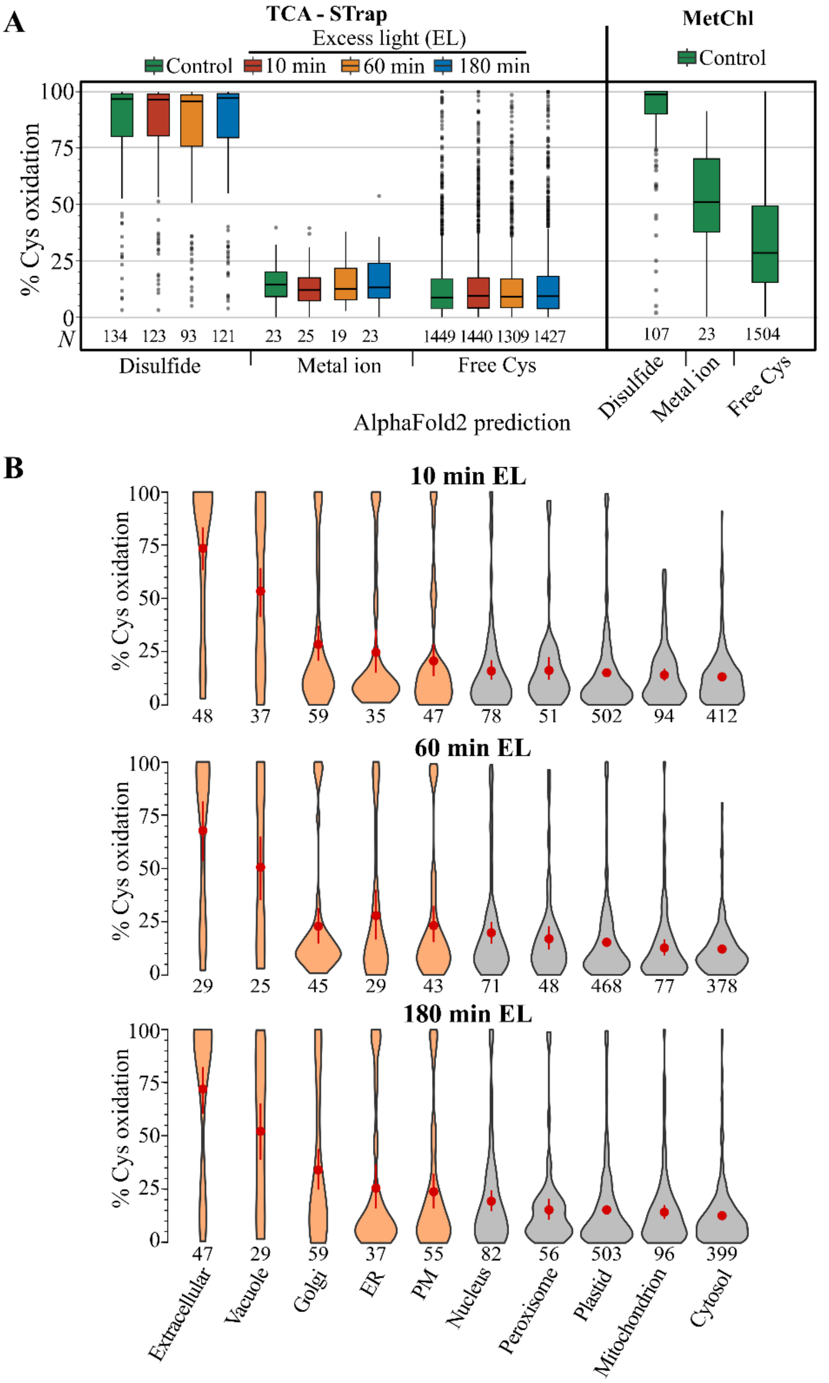


**Fig. S6. Cys oxidation degrees measured following acidic or non-acidic protein extraction.**

(**A**) Cys oxidation degrees quantified in three out of four replicates categorized according the AlphaFold2-based prediction as part of a disulfide, metal ion-binding pocket or free thiol (*33*). Next to the results obtained after trichloroacetic acid – suspension trapping (TCA-Strap, *left panel*), the Cys oxidation distribution after a non-acidic methanol-chloroform (MetChl) protein extraction is shown (*right panel*). (**B**) Oxidation degree for Cys quantified (after TCA-STrap) in three out of four replicates in the EL conditions according subcellular locations predicted by the SUBAcon algorithm (*75, 76*). Compartments associated with the secretory pathway are colored orange, with the mean oxidation degree ± sd in red and the number of Cys peptide precursors indicated.

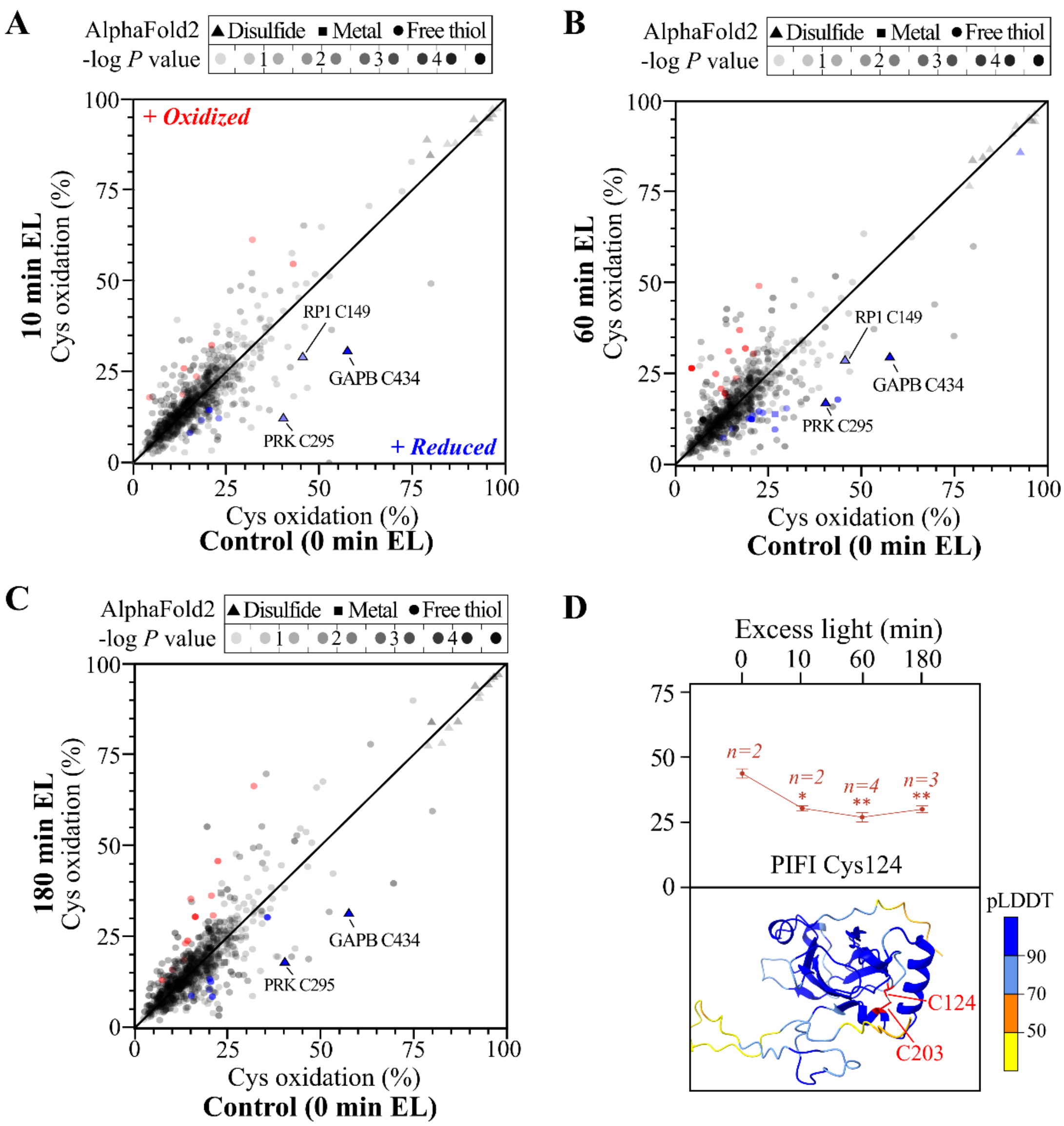


**Fig. S7. Differential cysteine oxidation degrees upon excess light treatment as measured by MaxQuant labeled analysis.**

(**A-C**) Scatterplot of the degree of Cys oxidation in unstressed plant leaves (control, x-axis) and leaves exposed to 10 (**A**), 60 (**B**) or 180 minutes of EL (**C**) (y-axis). Cys transparency is according the *P* value of the Welch's t-test (EL / control), with significantly altered oxidation ($P < .05$, $\geq 5\%$ absolute change) indicated in red (more oxidized) or blue (more reduced). Shapes were according to the AlphaFold2 predicted annotation (*33*). (**D**) Degree of oxidation for Cys124 of POST-ILLUMINATION CHLOROPHYLL FLUORESCENCE INCREASE (PIFI). The number of replicates (*n*) in which an oxidation degree was quantified is shown and asterisks indicate *P* values of the Welch's t-test comparing the excess light and control condition oxidation degrees (* < 0.05, ** < 0.01, and *** < 0.001). Protein structure was predicted by AlphaFold2 (*51*). The Cys124-203 disulfide was indicated in red, and residues were colored according the per-residue confidence score (pLDDT) with >90 being "highly confident" (blue), pLDDT > 70 being "low confident" predictions (light blue), and yellow and orange corresponding to lower confident predictions (pLDDT <70 and <50, respectively).

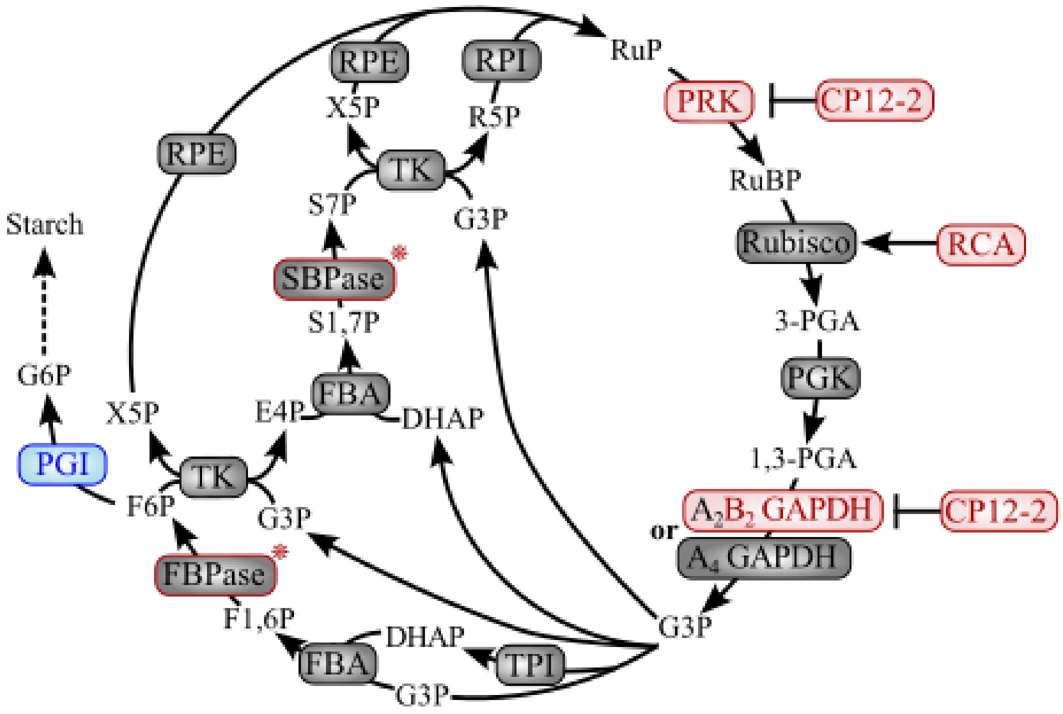


**Fig. S8. Redox regulation in the Calvin-Benson cycle and associated proteins.**

Characterized redox-regulated proteins are indicated in red, with those identified in this study as reduced under excess light in a red background and red text (PRK, CP12-2, RCA, and GAPB), while no peptides matching the active site of FBPase and SBPase were identified (asterisk). Plastidial PGI is proposed as a novel redox-regulated enzyme (blue). Pathway figure based on (*36*). Abbreviations: PGI, phosphoglucoisomerase (plastidic), PGK, phosphoglycerate kinase; GAPDH, glyceraldehyde-3-phosphate dehydrogenase; TPI, triose phosphate isomerase; FBA, fructose-1,6-bisphosphate aldolase; FBPase, fructose-1,6-bisphosphatase; TK, transketolase; SBPase, sedoheptulose-1,7-bisphosphatase; RPE, ribulose-5-phosphate 3-epimerase; RPI, ribose-5-phosphate isomerase; PRK, phosphoribulokinase. Metabolites, RuBP, ribulose-1,5-bisphosphate; 3-PGA, 3-phosphoglycerate; 1,3-PGA, 1,3-bisphosphoglycerate; G3P, glyceraldehyde-3-phosphate; DHAP, dihydroxyacetone phosphate; F1,6P, fructose-1,6-bisphosphate; F6P, fructose-6-phosphate; G6P, glucose-6-phosphate; X5P, xylulose-5-phosphate; E4P, erythrose-4-phosphate; S1,7P, sedoheptulose-1,7-bisphosphate; S7P, sedoheptulose-7-phosphate; R5P, ribulose-5-phosphate; RuP, ribulose-5-phosphate.

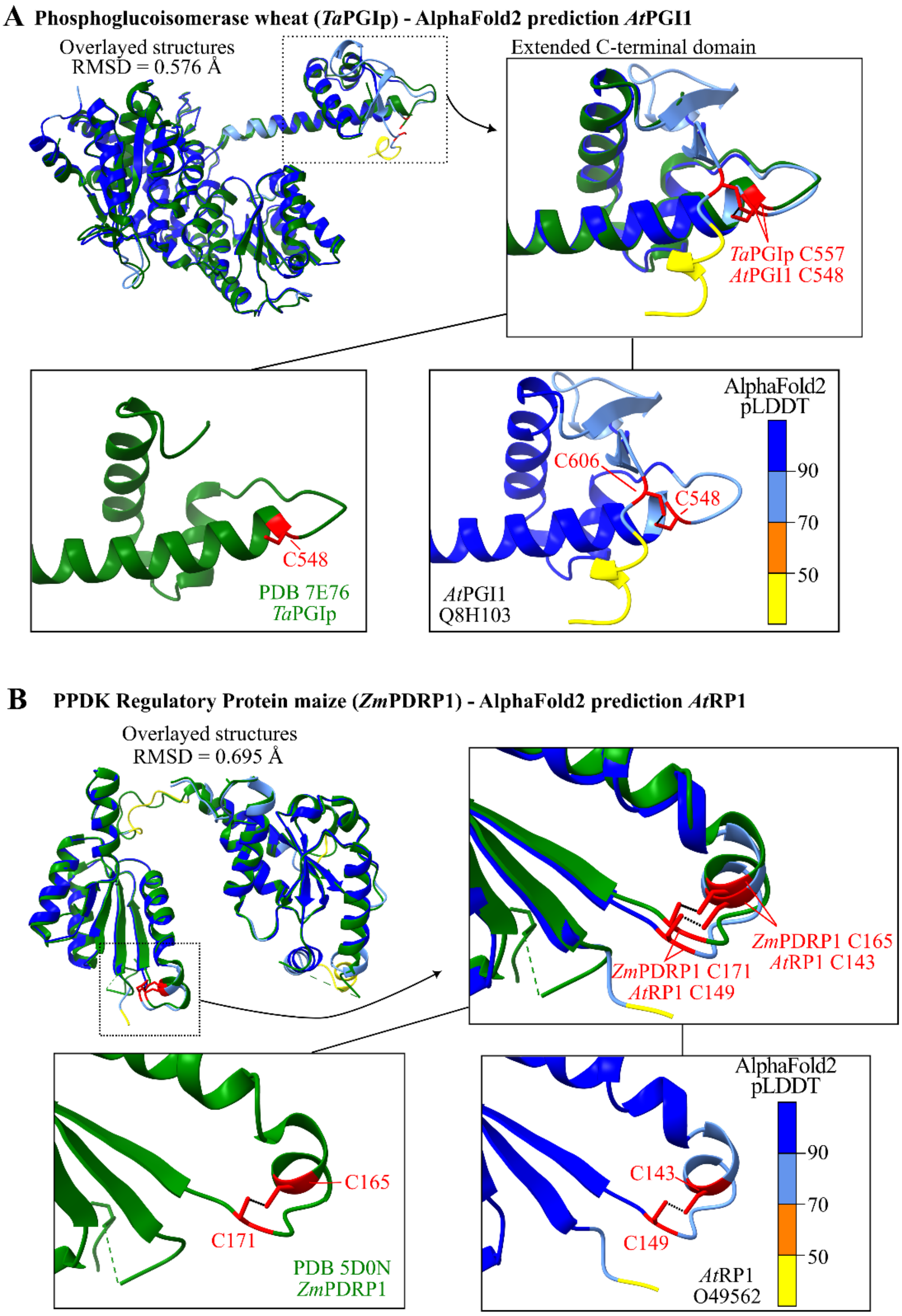


**Fig. S9. AlphaFold2 prediction comparisons against experimental crystal structures.**

(**A**) Comparison of a *Triticum aestivum* plastidic phosphoglucoisomerase (*Ta*PGIp) crystal structure (*67*) with the PGI1 predicted structure of Arabidopsis. The last 22 residues of TaPGIp were unmodeled, thus lacking the Cys corresponding to Cys606 in Arabidopsis. Inset boxes display the C-terminal extended domain containing a potential disulfide involved in redox regulation. (**B**) Comparison of *Zea mays* PPDK regulatory protein 1 (*Zm*PDRP1) crystal structure (*66*) with the RP1 predicted structure in Arabidopsis. Zoom-in boxes display a potential disulfide involved in redox regulation. For both panel cysteines part of (predicted) disulfides were colored red. Structures derived from crystal structures were colored green, while AlphaFold2 residues were colored according the per-residue confidence score (pLDDT) with > 90 being highly confident and < 50 being of very low confidence. Structures were superimposed using the built-in Matchmaker tool and visualized using ChimeraX (v1.6.1) (*77*).

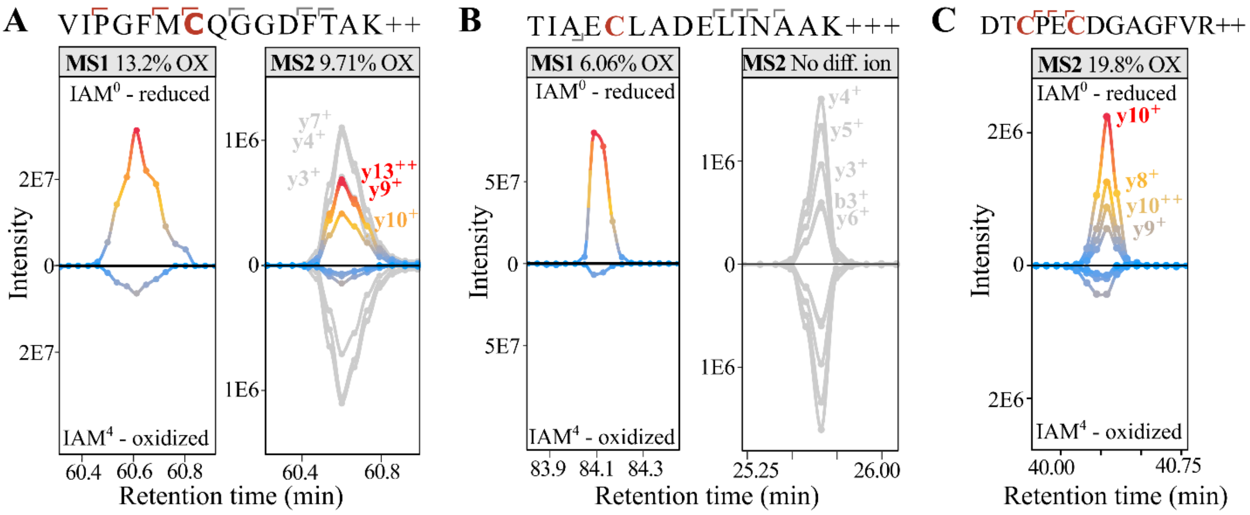


**Fig. S10. MS1 and MS2 level cysteine oxidation degree profiling.**

Extracted ion chromatogram peak areas at MS1 and MS level for cysteine-containing peptides labelled by $IAM^0$ (oxidized, top) and $IAM^4$ (reduced, bottom). Peaks were color-coded according their intensity. At MS2 level, grey lines indicate non-differentiating fragment ions (lacking Cys) and intensity-based coloured lines indicate differentiating ions (containing $IAM^4/IAM^0$ labelled Cys). Showing examples for peptides with a single Cys containing differentiating fragment ions (A) and lacking differentiating fragment ions (B). In addition, for peptides containing more than one Cys, differentiating fragment ions can point to the oxidation degree for specific Cys within these isobaric peptides, as illustrated with differentiating y-ions for the second Cys within ‘DTCPECDGAGFVR’ (C).

**Other Supplementary Materials**

**Data S1. Quantified cysteine oxidation degrees per sample and differential oxidation by excess light treatment. (separate file)**

Detailed legend see Excel file.

**Data S2. Quantified cysteine oxidation degrees per sample by plexDIA in methanol-chloroform extraction of unstressed leaves. (separate file)**

Detailed legend see Excel file.

**Data S3. Data Differential protein abundance analysis MSstats. (separate file)**

Detailed legend see Excel file.

**Data S4. Combined S-sulfinylation ($SO_2H$) and S-sulfonylation ($SO_3H$) sites identified in a label-free MaxQuant search. (separate file)**

Concatenated $SO_2H$ and $SO_3H$ modification site reports outputted by MaxQuant in a label-free search of the DDA data.
